# Stratum Lacunosum-moleculare Interneurons of the Hippocampus Coordinate Memory Encoding and Retrieval

**DOI:** 10.1101/2021.06.11.448103

**Authors:** Jun Guo, Heankel Cantu Oliveros, So Jung Oh, Bo Liang, Ying Li, Ege T. Kavalali, Da-Ting Lin, Wei Xu

## Abstract

Encoding and retrieval of memory are two processes serving distinct biological purposes but operating in highly overlapping brain circuits. It is unclear how the two processes are coordinated in the same brain regions, especially in the hippocampal CA1 region where the two processes converge at the cellular level. Here we find that the neuron-derived neurotrophic factor (NDNF)-positive interneurons at stratum lacunosum-moleculare (SLM) in CA1 play opposite roles in memory encoding and retrieval. These interneurons show high activities in learning and low activities in recall. Increasing their activity facilitates learning but impairs recall. They inhibit the entorhinal- but dis-inhibit the CA3- inputs to CA1 pyramidal cells and thereby either suppress or elevate CA1 pyramidal cells’ activity depending on animal’s behavioral states. Thus, by coordinating entorhinal- and CA3- dual inputs to CA1, these SLM interneurons are key to switching the hippocampus between encoding and retrieval modes.

## Introduction

Memory encoding and retrieval are two closely linked biological processes: during encoding, plastic alterations occur in the brain to store information, while during retrieval the stored information is reactivated so that we can re-experience the past. The accuracy of memories is predicted to rely on how closely the retrieval matches the encoding ^1^. Indeed, the key brain regions for retrieval are also critically engaged in memory encoding ^2–4^. Recent studies showed that neurons active in encoding are reactivated during retrieval ^5^; artificially activating these neurons leads to memory-guided behaviors ^6–8^, while silencing them impairs retrieval ^9^. Together, the findings show that encoding and retrieval overlap at systems, circuits and cellular levels.

Through operating in the same brain circuits, encoding and retrieval are two distinct processes fulfilling different needs. During encoding, the plasticity of neuronal network is essential to writing and storing information^10^. Conversely, during retrieval, plasticity may compromise the stored memories. Although memories are indeed dynamic and subjected to consolidation, reconsolidation and extinction, many memories are repeatedly retrieved without losing accuracy^11^. Furthermore, cognitive processing requires that neuronal activities unequivocally represent either ongoing or past events. Thus encoding and retrieval should be spatially or temporally separated to avoid mingling and interference ^12,13^. Asymmetry between the two processes at the brain systems level has been noted ^14^. A recent work also demonstrates that some hippocampal CA1 pyramidal cells may preferentially engage in either encoding or retrieval ^15^. However, how encoding and retrieval are coordinated and separated in overlapping brain circuits and neurons remains elusive.

The hippocampal CA1 region is involved in both encoding and retrieval^6,15–19^. CA1 principal cells, the pyramidal neurons, receive two major excitatory inputs from the entorhinal cortex layer 3 (EC3) and CA3, respectively ^20^. The EC3 inputs form synapses on the distal dendrites of the pyramidal neurons in stratum lacunosum-moleculare (SLM); while the CA3 inputs terminate on the proximal dendrites in stratum radiatum (SR) and stratum oriens (SO). The synaptic activities of the EC3 and CA3 inputs reach CA1 sequentially in theta cycles ^21–25^. Hasselmo and colleagues proposed that encoding and retrieval are segregated to different theta phases to minimize their interference ^13^. In this model, encoding occurs when the EC3 inputs are strong but CA3 inputs are weak. Simultaneously, CA3 inputs have strong long-term potentiation (LTP). On the contrary, retrieval occurs when EC3 inputs are weak but CA3 inputs are strong. Consistent with this hypothesis, silencing CA1 neuronal activity at specific phases of theta selectively facilitates encoding or retrieval^26^. This model is also supported by increasing evidence that the dual inputs differentially contribute to encoding and retrieval^12,22–25,27^. However, it is not clear why synaptic potentiation preferentially occurs when synaptic transmission is weak. In fact, a low level of synaptic activity may not favor synaptic potentiation due to a lack of “cooperativity” among synapses ^28^.

The CA1 dual inputs and the corresponding pyramidal cell dendritic compartments are regulated by GABAergic interneurons ^29–34^. These inhibitory neurons may hold the key to solve this paradox. Many interneurons are embedded among the EC3 inputs at SLM. The physiological properties of these interneurons have been carefully examined ^36,37,33,35^, but their contribution to learning and memory remains unknown. In this study, we selectively targeted, monitored and controlled a major fraction of the SLM interneurons. We find that these interneurons coordinate memory encoding and retrieval by differentially regulating the dual inputs to CA1. The functions of these interneurons in local circuits may reconcile the need of synaptic potentiation and the simultaneous weak synaptic transmission during encoding.

## Results

### 1. Expression of NDNF at SLM of the CA1

Neuron-derived neurotrophic factor (NDNF) is a genetic marker for a subset of GABAergic neurons, especially neurogliaform cells in the cortex ^36,37^. *In situ* hybridization of NDNF (Allen brain atlas) demonstrated that in the hippocampus NDNF was selectively expressed at SLM. To further determine the expression pattern of NDNF in the hippocampus, we used RNAscope *in situ* hybridization to assay NDNF along with a few selected markers for GABAgeric interneurons in the hippocampus, including Glutamate decarboxylase 1 (GAD1), Cholecystokinin (CCK), Neuropeptide Y (NPY) and Nitric Oxide Synthase 1 (NOS1) (Fig. 1). Since some markers showed an enrichment at the border between SR and SLM, presumably stratum lacunosum (S-L), we counted the labeled cells in S-L (defined as within 25 μm from the SR-SLM border) and stratum moleculare (S-M, the rest of SLM) separately. The pan-interneuron marker GAD1 labeled neurons in all layers of the CA1 (Fig. 1a). At SLM where GABAergic interneurons were the only neuronal type, GAD1 labeled cells with large sizes (>10 μm in their shorter axis), indicating that these were the sizes of neuronal soma at SLM. NDNF-positive cells were observed in both large and small (<10 μm in shorter axis) sizes, indicating that NDNF was expressed by both neuronal and non-neuronal cells at SLM. Therefore, we only counted NDNF-positive cells with more than 10 μm in their shorter axis as neurons. 95% of the NDNF-positive neurons (referred to as NDNF-cells) expressed GAD1 and accounted for 53% of the GAD1-positve neurons at SLM, with a higher percentage at S-M (72%) than that at S-L (35%) (Fig. 1a, 1b, 1e). This showed that NDNF identified a major fraction of SLM interneurons. 78% of NDNF-positive neurons expressed NPY, a marker for neurogliaform cells (Fig. 1b, 1c, 1e) ^38^. NDNF showed minimal overlap with CCK, which was enriched in SL. Both caudal ganglionic eminence (CGE) and medial ganglionic eminence (MGE)-derived neurogliaform cells are present in SLM ^39^. NOS1 labels interneurons derived from MGE. Consistent with recent single cell genetic profiling of hippocampal interneurons ^40^, NOS1 was expressed in only 10% of NDNF-cells (Fig. 1b, 1d, 1e), indicating that most NDNF cells come from CGE. Together, the results indicate that NDNF is expressed by a major fraction of GABAergic neurons at SLM, and the majority of them express the marker of neurogliaform cells.

**Figure 1.**
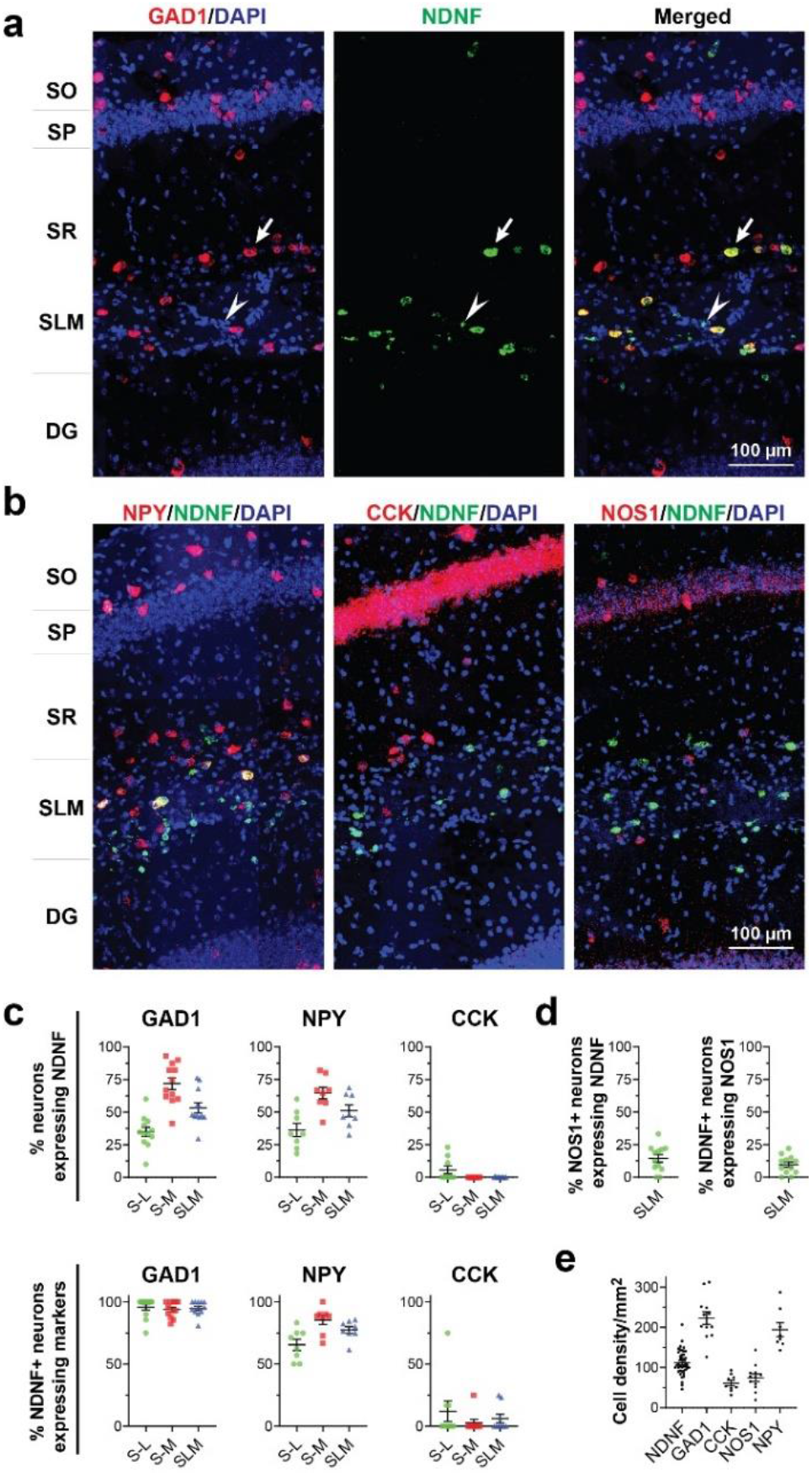
Expression of NDNF by interneurons at SLM of the CA1. (**a**) Representative RNAscope *in situ* hybridization photos showing the co-localization of GAD-1 and NDNF. The arrow and arrowhead indicate cells with distinct soma sizes. (**b**) Representative photos showing the co-localization of NDNF with NPY, CCK or NOS1, respectively. (**c**) Quantification of the co-localization of NDNF with GAD-1, NPY, and CCK, respectively. (**d**) Quantification of the co-localization of NDNF with NOS1. (**e**) The cell density of SLM interneurons expressing NDNF, GAD1, CCK, NOS1 and NPY. Abbreviations: SO, stratum oriens; SP, stratum pyramidale; SR, stratum radiatum; S-L, stratum lacunosum; S-M, stratum moleculare; SLM, stratum lacunosum-moleculare.

### 2. Selective targeting of NDNF-cells at SLM

To functionally characterize NDNF-cells, we tried to target these cells selectively with a Cre driver mouse line, Ndnf-IRES2-dgCre (referred to as NDNF-Cre mice from now on) (Fig. 1 and Fig. S1). This or similar mouse lines have been used to target NDNF-positive cells in the cortex ^36^ or hippocampus for electrophysiogy^41^. We first crossed Ndnf-IRES2-dgCre mice with tdTomato reporter mice (Ai9 mice) to check the distribution of Cre activity. tdTomato expression was detected in not only the expected interneurons but also multiple cell types across the brain, consistent with the RNAscope results above (Fig. S1a). To exclude the possibility that NDNF gene was transiently turned on during development, we injected into the CA1 of adult mice adeno-associated virus (AAV) which mediated Cre-dependent expression of fluorescent proteins under the control of synapsin promoter. We found that in addition to interneurons, Cre activity was present in some excitatory CA1 pyramidal neurons and DG granule cells in adult mouse brain (Fig. S1b). This leaky expression of Cre activity in excitatory neurons was unexpected since NDNF was not expressed in these cells (Fig. 1). However, it was consistent with the data obtained with Ai9 mice (Fig. S1a), suggesting that at the locus of NDNF gene Cre may be expressed at low level but it is sufficient to turn on reporter genes. Since this leakage was not reported in previous study using these mice ^41^, it could also be due to the instability of Cre-drive lines as frequently noted. To further improve the selectivity of targeting, we took an intersectional strategy by constructing an AAV vector AAV hDlx-DIO-EGFP to mediate Cre-dependent expression of EGFP under a synthetic hDlx promoter known to mediate GABAergic neuron-specific expression (“DIO” represents “double-floxed inverted open reading frame”) ^42^. After injecting this AAV into the CA1, we observed EGFP-positive cells solely at SLM, which had only GABAergic interneurons but not excitatory neurons, indicating selective targeting of the NDNF-positive interneurons at SLM (referred to as NDNF-cells) (Fig. 2a).

**Figure 2.**
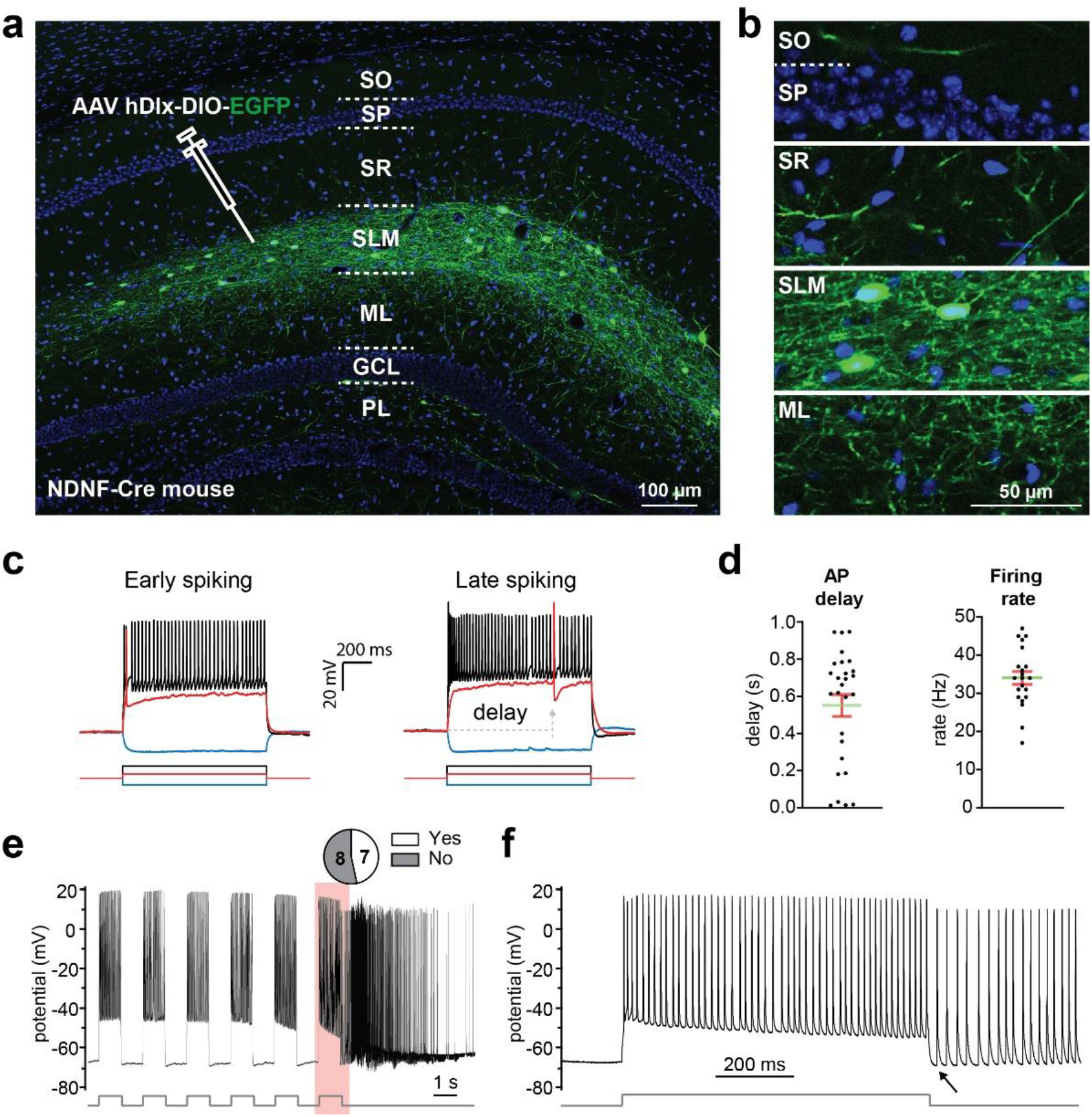
Selective targeting of neurogliaform cells at SLM. (**a, b**) Interneurons at SLM selectively expressed EGFP. AAV hDlx-DIO-EGFP was injected into the CA1 of NDNF-Cre mice. Abbreviations: SO, stratum oriens; SP, stratum pyramidale; SR, stratum radiatum; SLM, stratum lacunosum-moleculare; ML, molecular layer; GCL, granule cell layer; PL, polymorphic layer. The blue channel is counterstain with DAPI. (**b**) EGFP-positive axonal fibers of NDNF-cells were distributed at multiple layers of the hippocampus. (**c, d**) Representative traces and quantification of electrophysiological properties of early spiking and late spiking NDNF-cells. The bars are means ± SEM (error bars); (**e, f**) Repetitive depolarization triggered persistent firing. (**f**) Enlargement of the pink color-shaded area in **e**. The arrow indicates the start of spontaneous persistent firing. “Yes/No” indicates if persistent firing was triggered.

NDNF-cells had dense plexus of neurites at SLM (Fig. 2a). Their axons were concentrated at SLM and sparsely extending to other layers at CA1, DG and sparsely at the retrosplenial cortex (RSP) (Fig. 2b). Whole-cell patch clamp recordings demonstrated that the majority of NDNF-cells were late firing (Fig. 2c, 2d). Furthermore, repetitive current injections triggered persistent firing in a significant fraction of NDNF-cells (7 out of 15 cells) (Fig. 2e, 2f), suggesting that the majority of NDNF-cells were neurogliaform cells ^33,38^.

### 3. Monosynaptic inputs to NDNF-cells

To determine the synaptic inputs to NDNF-cells, we conducted retrograde transneuronal tracing with genetically modified rabies virus--the RabiesΔG (EnvA) system ^43^. We expressed in NDNF-cells the receptor for RabiesΔG(EnvA)-mCherry, TVA, and optimized rabies glycoprotein (oG), (Fig. 3a, 3b). We detected mCherry-positive neurons—mono-synaptic presynaptic neurons to NDNF-cells--in multiple brain regions but mostly in the entorhinal cortex, CA3, and CA1, subiculum and medial septum (Fig. 3b-d, Fig. S2a). These results indicate that NDNF-cells are mainly innervated by the entorhinal cortex, but they may also integrate information from broadly distributed neurons. The thalamus, especially the nucleus reuniens, also projects to CA1 and form synapses at SLM. Slice electrophysiology study has shown that interneurons at SLM were innervated by the thalamic inputs ^44^. In our tracing, we only detected small number of neurons at nucleus reuniens (Fig. 3d). This may be because that thalamic inputs are more concentrated in the ventral part of the hippocampus while our viral injections were at the dorsal part^45^.

**Figure 3.**
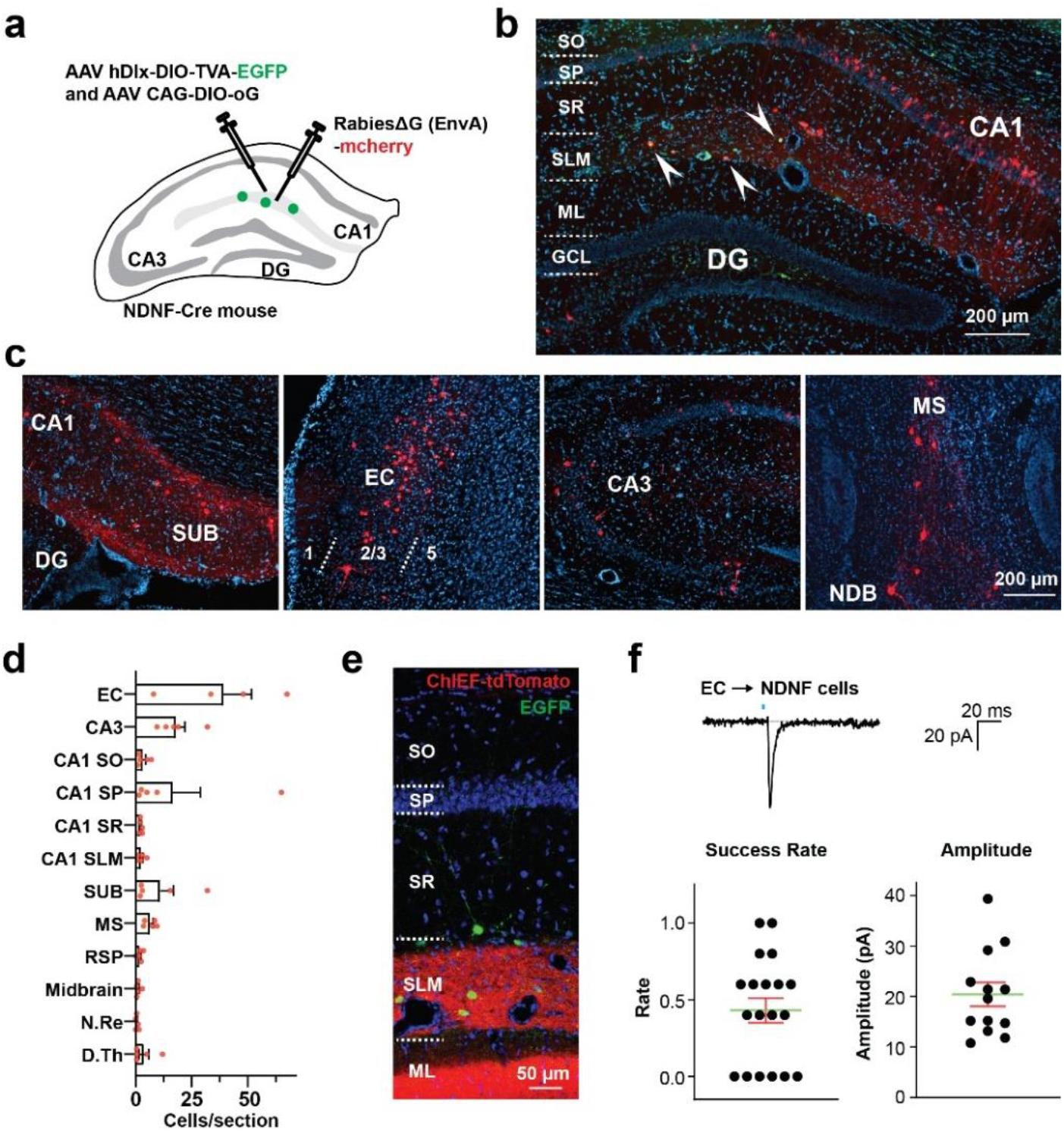
Monosynaptic inputs to NDNF-cells. (**a**) Scheme of the experimental procedure for retrograde transneuronal tracing of NDNF-cells with recombinant rabies virus. (**b**) Representative photo showing the starter neurons at SLM (dual labeled by EGFP and mCherry) and the retrogradely traced neurons at CA1 (labeled by mCherry). The blue channel is counter staining with DAPI. (**c**) Distribution of retrogradely traced neurons in the subiculum (SUB), entorhinal cortex (EC), CA3, medial septum (MS) and nucleus of the diagonal band of Broca (NDB). (**d**) Quantification of the distribution of the retrogradely traced neurons (n=5 mice). (**e-f**) Electrophysiology verification of the synaptic inputs to NDNF-cells. (**e**) Representative photo showing ChIEF-tdTomato-positive axonal terminals from EC and the EGFP-positive NDNF-cells at SLM. (**f**) Representative traces and quantification of EPSCs recorded in NDNF-cells. EPSCs were evoked by optical stimulation applied to EC-derived axonal terminals. Success rate is the ratio of responsive trials in 5 consecutive trials. The bars are means ± SEM.

To verify the synaptic connectivity, we expressed channelrhodopsin ChIEF in EC and recorded EGFP-positive NDNF-cells at SLM (Fig. 3e, Fig. S2b). As expected, photostimulation evoked excitatory postsynaptic currents (EPSCs) in the majority of the recorded cells, confirming functional synapses between entorhinal neurons and NDNF-cells (Fig. 3f).

### 4. Opposite roles of NDNF-cells in learning and recall

To examine if and how NDNF-cells contribute to learning and memory, we made AAV hDlx-DIO-hM3Dq-mCherry to selectively express in NDNF-cells the Designer Receptors Exclusively Activated by Designer Drugs (DREADD) effector protein hM3Dq, which activates neurons in the presence of low concentrations of clozapine ^46^ (Fig. 4a). AAV hDlx-DIO-hM3Dq-mCherry was injected to the CA1 of NDNF-Cre mice (Cre+) or their Cre- littermate controls. Then these mice underwent contextual fear conditioning. In the fear conditioning training, after exploring the conditioning chamber for 2 minutes, the mice received a 30-second tone that co-terminated with a 2-second footshock (Fig. 4c). The single-shock fear conditioning has been showed to consistently generated hippocampus-dependent memory^47^. We administered clozapine 30 min before training so that NDNF-cells were activated during training. The mice were tested in the original training context on the 2nd day (Context test) without tone; and tested in an altered context--a new chamber sharing some features with the training context but containing features distinct from the original context--on the 3rd day for 5 minutes (Altered context test). After 5 min of altered context test, the mice were present with the tone for 1 minute in the altered context (Tone test). The Cre+ mice recalled significantly better than their littermate controls in the context test (Fig. 4d). The freezing to the altered context or tone was not significantly different between the Cre− and Cre+ mice (Fig. S3a).

**Figure 4.**
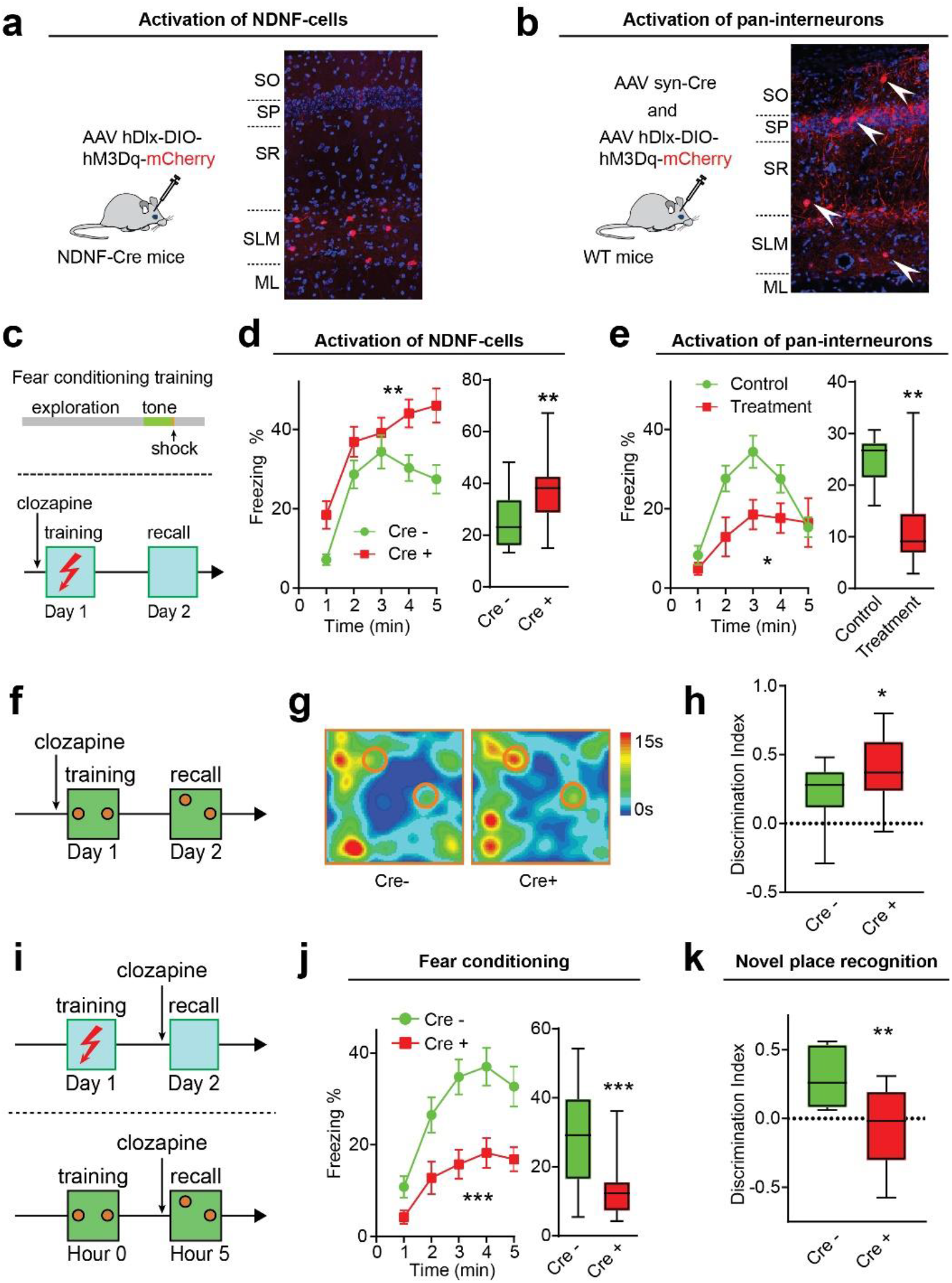
Opposite roles of NDNF-cells in learning and recall. (**a**) hM3Dq-mCherry was expressed exclusively in NDNF-cells at SLM in NDNF-Cre+ mice. (**b**) hM3Dq-mCherry was expressed in pan-interneurons at CA1 in wild-type mice. (**c**-**e**) Fear conditioning when NDNF-cells were activated. Mice were trained with a single tone-footshock pair (**c**, upper panel) and received clozapine (0.1 mg/kg, i.p.) 30 min before training to activate hM3Dq (**c**, lower panel). (**d**) Pre-training administration of clozapine in Cre+ mice increased their freezing level in recall. n=16-17 mice in each group. (**e**) Pre-training administration of clozapine to activate pan-interneurons in CA1 in wild-type mice decreased their freezing level in recall. n=10 mice in each group. (**f**-**h**) Novel place recognition with pre-training administration of clozapine. (**g** and **h**) Representative heat-maps of mice travelling locations (**g**) and the discrimination index of new versus old locations [discrimination index = (Time_new_–Time_old_) / (Time_new_+Time_old_, **h**). Pre-training administration of clozapine improved the performance of NDNF-Cre+ mice. n=16-17 mice in each group. (**i**-**k**) Fear conditioning and novel place recognition with pre-recall administration of clozapine. (**j**) Pre-recall administration of clozapine reduced freezing in NDNF-Cre+ mice in context test of fear conditioning. n=17-20 mice in each group. (**k**) Pre-recall administration of clozapine impaired the performance of NDNF-Cre+ mice in novel place recognition test. n=9 mice in each group. Data are means ± SEM; statistical significance (*P < 0.05; **P < 0.01, ***P < 0.001) was assessed by two-way ANOVA (for data in line graphs) or Student’s t-test (column graphs).

We also tested how global activation of CA1 interneurons affect learning and memory (Fig. 4b). The treatment group were wild-type mice that received injection of AAV hDlx-DIO-hM3Dq-mCherry with AAV expressing Cre (AAV Syn-Cre) to express hM3Dq in broadly distributed interneurons at CA1. The control group were mice injected with AAV hDlx-DIO-hM3Dq-mCherry but not AAV Syn-Cre. When tested with the same fear conditioning protocol with clozapine injected 30 min before training, the treatment group did worse in the context test (Fig. 4e), demonstrating that non-specific activation of CA1 interneurons impaired learning. The results indicate that contrary to global silencing of CA1, selective activation of NDNF-cells enhances learning.

To check if activation of NDNF-cells improves fear conditioning learning only, we tested another hippocampus-dependent task--novel place recognition test. NDNF-Cre+ mice performed better than the controls when they received clozapine before training (Fig. 4f-h, Fig. S4a). The experiments demonstrate that increasing NDNF-cells activity facilitates hippocampus-dependent learning.

We then tested if NDNF-cells affected memory retrieval. NDNF-Cre+ mice expressing hM3Dq in NDNF-cells and their littermates underwent fear conditioning training or novel place recognition training without clozapine, but received clozapine 30 min before recall tests. NDNF Cre+ mice performed significantly worse in both tests, indicating that activation of NDNF-cells impaired recall (Fig. 4i-k, Fig. S3b, Fig. S4a). Together, the data demonstrate that NDNF-cells at SLM exert opposite effects on memory encoding and retrieval.

We also inhibited NDNF-cells with pharmacogenetics. We made AAV hDlx-DIO-hM4Di-EGFP to express the inhibitory DREADD effector hM4Di in NDNF-cells. Unfortunately, hM4Di-EGFP was expressed at very low level at SLM (Fig. S5a) and we did not observe significant difference in fear conditioning between Cre− and Cre+ mice when clozapine was administrated 30 min before training (Fig. S5b). Further optimization of the AAV will be necessary for a conclusive experiment. We further tried optogenetic silencing of NDNF-cells. We constructed AAV hDlx-DIO-JAWS-EGFP to selectively express JAWS, an inhibitory opsin^48^, in NDNF-cells for temporally precise inhibition of NDNF-cells. JAWS was expressed at a high level at SLM (Fig. S5c). Light effectively silenced NDNF-cells in brain slices (Fig. S5d). However, when we used this AAV to silence NDNF-cell in the duration of fear conditioning training, we did not observe a significant effect on mouse behaviors (Fig. S5e). This lack of impact may be due to the spatial restriction of optical stimulation—light could only be delivered with sufficient intensity to the neurons adjacent to the tip of optic fibers. Since NDNF-cells only accounted for ∼ 50% of interneurons at SLM, it was also possible that the lack of effect from pharmacogenetic or optogenetic silencing was because of the compensation from the other interneurons at SLM.

We also examined if activation of NDNF-cells regulated the consolidation of memory in the post-training period. NDNF-Cre+ mice expressing hM3Dq in NDNF-cells and their littermate controls received clozapine right after fear conditioning training (Fig. S4b). We found no significant difference between Cre+ and Cre− mice. We also examined the mice in open-field and elevated plus maze with clozapine administered, and observed no significant changes, suggesting the effects were specific to memory functions (Fig. S4c-d).

### 5. Distinct activity levels of NDNF-cells in learning and recall

To understand the roles of NDNF-cells in learning and memory, we monitored their activity by calcium imaging with the calcium indicator jGCaMP7f expressed in NDNF-cells. We implanted above SLM a GRIN lens to image NDNF-cells in awake mice with a head-mounted minimized microscope (Fig. 5a, Fig. S6a). Due to the low density of NDNF-cells and the short work distance of the GRIN lens, 2-5 cells were detected in each mouse (Fig. 5b, Movie S1). After motion correction, the calcium signals of individual neurons were extracted and plotted. NDNF-cells showed diverse kinetics in calcium signals (Fig. 5c, Fig. S6a). Due to the difficulty in reducing the calcium transients into individual spikes, we calculated the areas under the calcium traces to quantify the activity level of NDNF-cells (Fig. S6a). We recorded the mice in their home cages (Homecage) on day 0, during fear conditioning training (Training) on day 1, in context test on day 2, and in altered context and tone test on day 3 (Fig. 5c).

**Figure 5.**
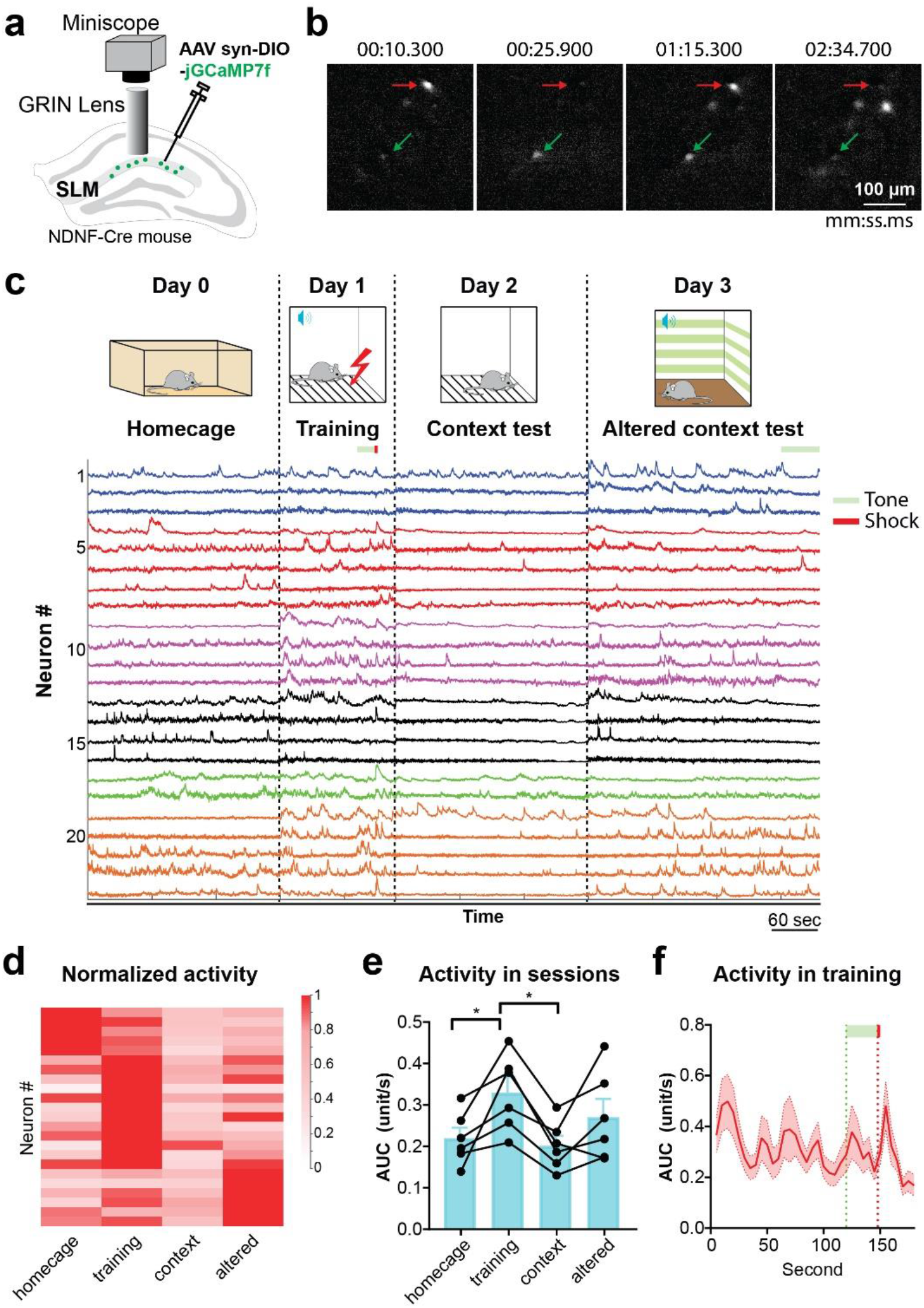
Distinct activity levels of NDNF-cells in learning and recall. (**a**) jGCaMP7f was selectively expressed in NDNF-cells, and GRIN lens was implanted above SLM for imaging NDNF-cells in behaving mice. (**b**) Representative photo frames of NDNF-cells imaged at different time points. The red or green arrows point to the same cell in the different frames. (**c**) Time course of 4 recording sessions (top) and the raw calcium signal traces of individual NDNF-cells. The mice underwent fear conditioning training and context test, followed by a session in an altered context. The traces from the same mouse are coded in the same color (n=6 mice). (**d**) Normalized activity of individual NDNF-cells in each of the recording session. Cells are sorted by their highest average activity in the sessions. (**e**) Plot of averaged calcium activities from each mouse in each of the 4 sessions (quantified by measuring the area under the calcium traces). The columns and bars are means ± SEM (n=6 mice); statistical significance (*P < 0.05) between each two conditions was assessed by paired t-test. One-way ANOVA of the 4 groups shows a significance difference between the conditions (p<0.05). (**f**) Binned neuronal activity of NDNF-cells during fear conditioning training. The data are the average of 23 neurons from 6 mice.

When mice were in their home cages, NDNF-cells showed basal activity (Fig. 5c-e). The activity increased in fear conditioning training, reaching a peak at the beginning of training, then gradually reduced, and finally increased again at the tone and footshock (Fig. 5f, the data did not reach statistical significance with the current sample size). In the context test (recall), NDNF-cells’ activity was low. When the mice were in an altered context with new features, the activity of NDNF-cells increased again (Fig. 5c-e). These findings suggest that NDNF-cells’ activity increases in response to learning (encoding) but decreases during retrieval.

### 6. State-dependent regulation of pyramidal neurons by NDNF-cells

To investigate how NDNF-cells differentially regulate encoding and retrieval, we imaged neuronal activities at the pyramidal layer. Calcium indicator GCaMP6s was expressed in the pyramidal layer and hM3Dq was expressed in NDNF-cells (Fig. 6a, 6b). GRIN lens was implanted right on top of the pyramidal layer. Due to the higher density of neurons in this layer, 64-163 cells were detected in each mouse (Fig. 6b, Movie S2). Since pyramidal layer consist of predominantly excitatory pyramidal neurons (over 90%) and a small interneuron population ^49^, the majority of recorded neurons were presumably pyramidal cells. When imaged in home cages, pyramidal neuronal activity in NDNF-Cre+ mice decreased 10 min after clozapine administration and remained decreased for more than 1 hour. However, in NDNF-Cre− mice administered with clozapine, or NDNF-Cre+ administered with saline and NDNF-Cre− mice administered with saline, pyramidal neuronal activity showed a minor reduction during the same period, possibly because mice grew accustomed to the imaging and reduced locomotion (Fig. 6c-e). The results show that activation of NDNF-cells inhibited pyramidal cells when the mice were in home cages, consistent with NDNF-cells’ identity as inhibitory interneurons. This effect was similar to that resulting from global activation of CA1 interneurons (Fig. S7).

**Figure 6.**
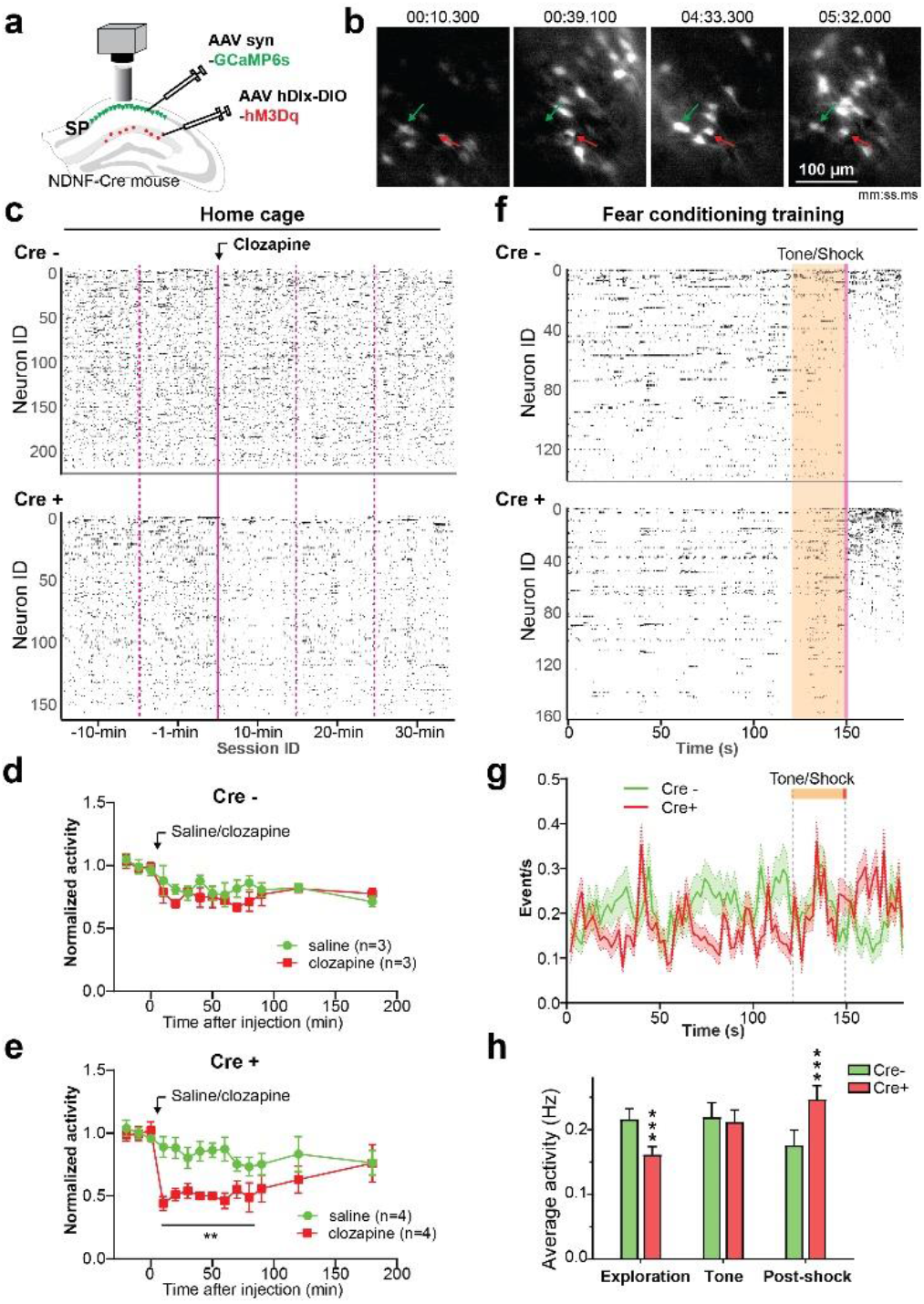
State-dependent regulation of pyramidal neurons by NDNF-cells. (**a**) Imaging calcium activity of neurons in CA1 pyramidal layer (SP) with hM3Dq selectively expressed in NDNF-cells. (**b**) Representative photo frames of neuronal calcium signals in the pyramidal layer. (**c** to **e**) Mice were imaged in 1-min sessions in the home cage. Clozapine or saline was administered right after the 1-min session. (**c**) Raster plots of neuronal calcium activity in a NDNF-Cre− and a Cre+ mouse. (**d** and **e**) Normalized average neuronal calcium activity before and after the administration of clozapine or saline in Cre− (**d**) or Cre+ (**e**) mice. Data are means ± SEM; statistical significance (**P < 0.01) was assessed by two-way ANOVA. (**f**) Representative raster plots of CA1 neuronal calcium activity of a NDNF-Cre− and Cre+ mouse in fear conditioning training. The neurons were sorted by their activity levels in the post-shock period. (**g**) CA1 neuronal activity from Cre+ or Cre− mice are averaged in 2s time bins. (**h**) Average CA1 neuronal activity of Cre+ and Cre− mice during the 2-min exploration before the tone, the tone, and the post-shock periods. Data are means ± SEM; statistical significance (***P < 0.001) was assessed by Mann-Whitney test.

We administered clozapine to the mice and imaged them during fear conditioning training. Neuronal activity in the pyramidal layer in NDNF-Cre+ mice was significantly lower in the first 2 min of exploration, compared to NDNF-Cre− control mice, similar to that in home cage. This difference largely disappeared in the 30 seconds of auditory cue. Surprisingly, after the foot shock, neuronal activity in NDNF-Cre+ mice increased (Fig. 6f-h). The results indicate the overall effect of NDNF-cells on CA1 pyramidal neurons depends on the animal behavioral states—it can be inhibitory or paradoxically excitatory.

### 7. Opposite regulation of the CA1 dual inputs by NDNF-cells

To understand how NDNF-cells produce opposite effects on pyramidal neurons, we analyzed the roles of NDNF-cells in local circuits with brain slice electrophysiology. We expressed opsin ChIEF in NDNF-cells with AAV hDlx-DIO-ChIEF-EGFP (Fig. 7a). 2-ms blue light pulses reliably triggered action potentials in NDNF-cells (Fig. S8a). Then we optically stimulated NDNF-cells and conducted extracellular recording at SLM, SR and SP, respectively. Activating NDNF-cells induced inhibitory postsynaptic field potential (fIPSP) selectively at SLM, consistent with their dense axonal terminals there (Fig. 7b and c). We also conducted whole-cell recording of the CA1 pyramidal cells and detected IPSPs or IPSCs induced by photostimulation of NDNF-cells (Fig. S8b-f). The IPSPs or IPSCs frequently showed a fast and a slow phase consisting of the GABA_A_ and GABA_B_ receptor-mediated inhibitory synaptic currents, respectively, which were well characterized in SLM interneurons ^33,38^. Antagonist of GABA_B_ receptor--CGP55845--selectively, blocked the slow phase. CGP55845 also increased the amplitude of the fast phase, indicating the NDNF-cells were subjected to tonic presynaptic inhibition mediated by GABA_B_ receptors (Fig. S8e). The IPSCs induced by stimulation of NDNF-cells showed a pronounced desensitization in a train of stimuli, similar to what has been described for some SLM interneurons^50^. These results indicate that NDNF-cells locally inhibit the distal dendrites of CA1 pyramidal neurons.

**Figure 7.**
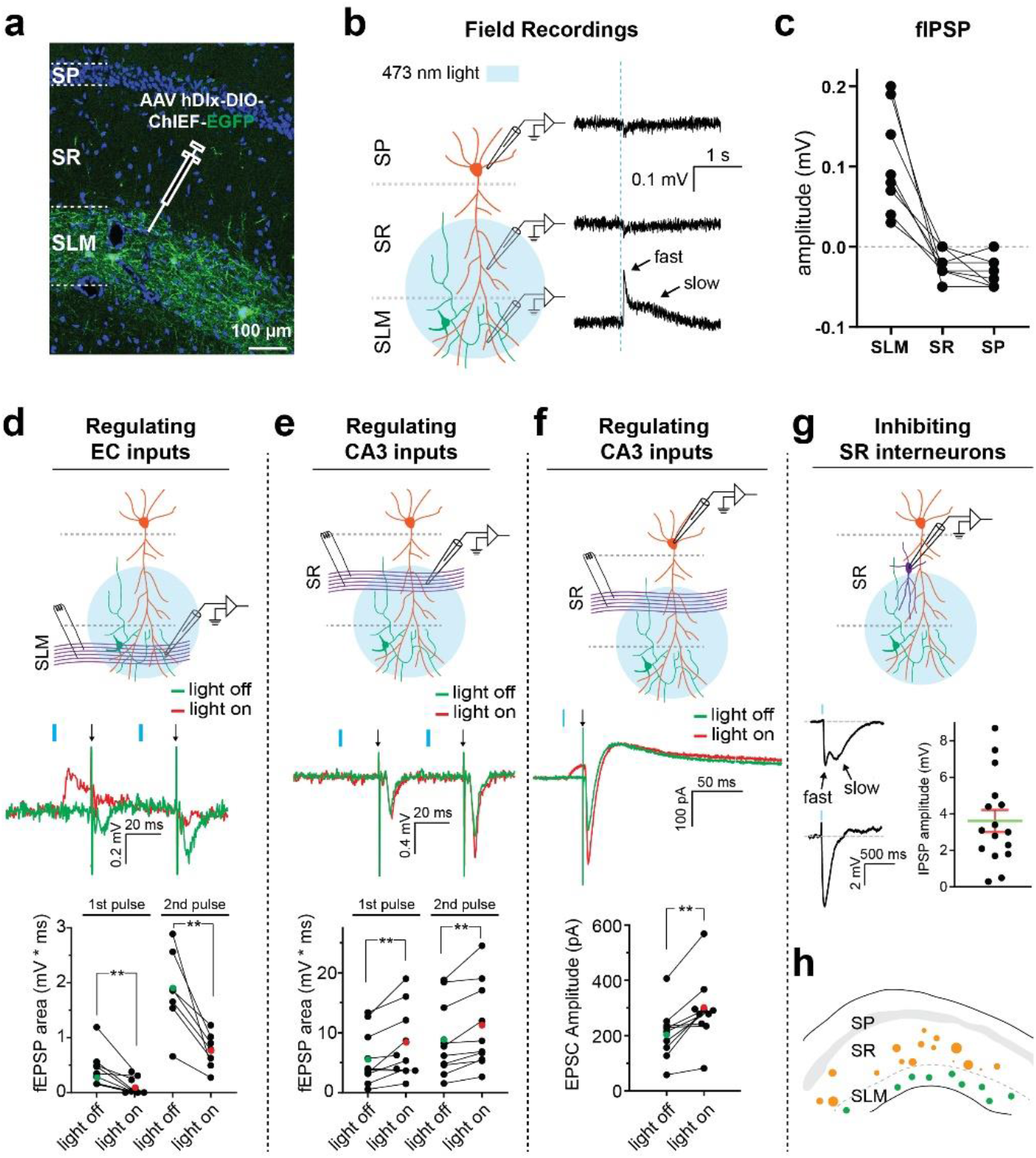
Opposite regulation of the dual inputs to CA1 by NDNF-cells. (**a**) Selective expression of ChIEF-EGFP in NDNF-cells. (**b**, **c**) Local field potentials were recorded at SLM, SR and SP, respectively, with photostimulation to NDNF-cells. (**d**) Field EPSPs evoked by electric stimuli applied to EC3-CA1 pathway, with (“light on”) or without (“light off”) photostimulation delivered to NDNF-cells 20 ms before the electric stimuli. The green and red dots are the means. Statistical significance between groups (**P < 0.01) was assessed by paired t-test. One-way ANOVA of the whole data set showed a significant difference (P<0.0001) between “light-on” and “light-off”. (**e**) Field EPSPs evoked by electric stimuli applied to CA3-CA1 pathway, with or without photostimulation to NDNF-cells. Statistical significance between groups (**P < 0.01) was assessed by paired t-test. One-way ANOVA of the whole data set showed a significant difference (P<0.0001) between “light-on” and “light-off”. (**f**) Whole-cell recordings of CA1 pyramidal cells. EPSPs were evoked by electric stimuli applied to CA3-CA1 pathway, with or without photostimulation to NDNF-cells. Statistical significance between groups (**P < 0.01) was assessed by paired t-test. (**g**) Whole-cell recordings of interneurons in SR with photostimulation applied to NDNF-cells. Photostimulation induced IPSP in all the cells recorded (n=16). The bars are means ± SEM. (**h**) Distribution of the recorded interneurons at SR. Each brown circle represents a recorded neuron with its area scaled to the amplitude of the IPSP recorded from the same cell.

To further analyze how NDNF-cells regulate EC3-to-CA1 and CA3-to-CA1synapses, we applied electric stimulation to either SLM or SR and conducted extracellular recordings at SLM or SR correspondingly (Although SLM also receives inputs from the thalamus, these thalamic inputs account for a small fraction especially in the dorsal portion of the hippocampus^45^). Since the activity of direct EC3-to-CA1 synapse is normally around 20 ms ahead of that of the indirect CA3-to-CA1 synapses ^21^, we delivered photostimulation of NDNF-cells 20 ms before applying electric stimulation to CA3-to-CA1 axons/synapses. For comparison, to EC3-to-CA1 axons/synapses we also delivered photostimulation 20 ms before the electric stimulation. Photostimulation of NDNF-cells reduced fEPSPs triggered by electric stimulation applied to SLM, consistent with our finding that NDNF-cells locally inhibit the distal dendrites of pyramidal cells (Fig. 7d). In contrast, photostimulation of NDNF-cells increased fEPSPs in SR (Fig. 7e). To further confirm that NDNF-cells facilitates SR excitatory synaptic activity, we conducted whole-cell recordings of the pyramidal neurons while photostimulating NDNF-cells. Consistent with above experiments, photostimulating NDNF-cells inducted IPSC in whole-cell recording, and increased the EPSC triggered by electric stimulation applied to synapses in SR (Fig. 7f), further confirming that NDNF-cells facilitate CA3-to-CA1synapses.

The facilitation of SR excitatory synaptic currents cannot be explained by direct inhibition of pyramidal cells. Instead, the facilitation may stem from dis-inhibition—inhibiting local inhibitory interneurons at SR, as reported in the hippocampus and other circuits ^51^. To test this possibility we conducted whole-cell recordings of SR interneurons. In all the SR interneurons recorded (15 cells from 4 mice), IPSPs were evoked by photostimulation of NDNF-cells, indicating that NDNF-cells dis-inhibited CA3-to-CA1 inputs (Fig. 7g, 7h). Some of the cells exhibited bi-phasic IPSPs, suggesting that the currents were mediated by both GABA_A_ and GABA_B_ receptors; in other neurons IPSPs showed only one phase, indicating that SR interneurons were heterogeneous. Together, the data show that NDNF-cells may dis-inhibit the excitatory synaptic activity through innervating SR interneurons.

### 8. Activity of CA1 dual inputs during learning

The inhibition and dis-inhibition indicate that NDNF-cells can adjust the gain of the dual inputs to CA1 pyramidal neurons. The local inhibition at the distal dendrites and dis-inhibition at proximal dendrites indicate that NDNF-cells can adjust the gain of the dual inputs and tune CA1 pyramidal neurons to one or the other. The net impact of NDNF-cells on pyramidal neuronal activity should be determined by the relative activities of the dual inputs to CA1 pyramidal neurons: when CA1 pyramidal neurons are driven primarily by EC3 inputs, activity of NDNF-cells inhibits the pyramidal cells; when CA1 pyramidal cells are driven mainly by CA3 inputs, NDNF-cells promote pyramidal neuronal activity. The increased neuronal activity at the pyramidal layer in fear conditioning training during the post-shock period caused by NDNF-cells activation (Fig. 6f-h) suggests that the pyramidal neurons are excited mainly by CA3 inputs in the post-shock period.

To test this possibility, we conducted calcium imaging of the synaptic terminals originating from EC3 and CA3, respectively. We expressed in EC3 or CA3 GCaMP6f fused with synaptic vesicle protein Synaptobrevin 2 (GCaMP6f-Syb2) (Fig. 8a, 8b), and implanted GRIN lens in CA1 above SLM or SR to image the collective calcium activity of the synaptic terminals (Fig. 8c). Global calcium signals fluctuated in both pathways in behaving mice (Fig. 8d, Movie S3). When calcium fluctuations were normalized to the average calcium activity in the first 2 minutes of fear conditioning training, CA3 inputs increased in the post-shock period (Fig. 8e). Together, these data indicate that NDNF-cells can both inhibit and dis-inhibit CA1 pyramidal neurons based on behavioral states of the mice.

**Figure 8.**
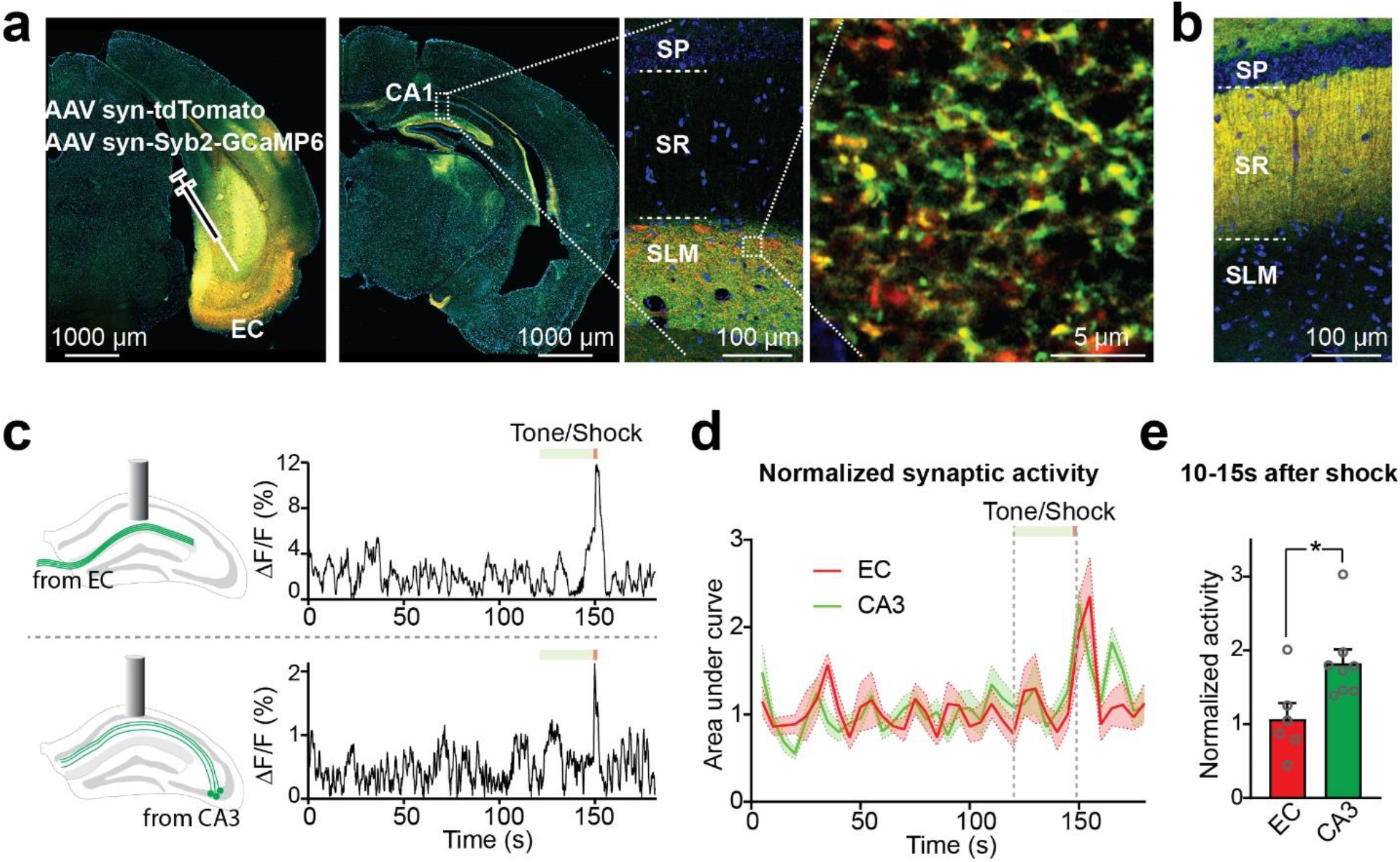
Activity of the dual inputs to CA1 during learning. (**a**) Targeted expression of Syb2-GCaMP6f at synaptic terminals in EC3-CA1 pathway. (**b**) Expression of Syb2-GCaMP6f in CA3-CA1 pathway. (**c**) Monitoring EC3-CA1 (top) or CA3-CA1 (bottom) synaptic activity via implanted GRIN lenses above either SLM or SR, respectively. Representative traces of the global calcium fluctuations of EC3-CA1 (top) or CA3-CA1 (bottom) synapses during fear conditioning training were shown. (**d**) Normalized activities of EC3-CA1 or CA3-CA1 synapses with 5-s bins. (**e**) Normalized activities of EC3-CA1 or CA3-CA1 synapses 15s after footshock. The circles represent data from each mouse. The column and bars are means ± SEM; statistical significance (*P < 0.05) was assessed by Mann-Whitney test (n=6-8 mice).

## Discussion

### 1. SLM NDNF-cells coordinate hippocampal information flow

SLM is home to a diversity of GABAergic interneurons. Their electrophysiological properties have been carefully examined over the years ^29,35^. Studies of their functional contributions to behavior have focused on the interneurons distributed at the border between SLM and SR. These border neurons are found to be innervated by entorhinal island cells and contribute to trace fear conditioning learning ^52^, or innervated by the long-range inhibitory inputs from the entorhinal cortex gating information flow into the hippocampus ^53^. They may also serve as a link between the thalamus and the retrosplenial cortex ^54^. In this study, we focused on the NDNF-positive interneurons that are distributed throughout SLM and account for about half of the total interneuron population there. The localization, expression of genetic markers and electrophysiological properties indicate that these neurons are mainly neurogliaform cells (Fig. 1-2).

SLM interneurons are shown to be primarily innervated by EC3 synapses and fire at the peak of theta oscillations ^38,55^, indicating that these neurons mediate feedforward inhibition in the EC3-to-CA1 pathway. Consistent with this, monosynaptic tracing showed that NDNF-cells were mainly innervated by the neurons in EC (Fig. 3). Besides EC, we also found their presynaptic neurons distributed in the CA3, CA1, subiculum, thalamus and many other regions. In CA1, the presynaptic neurons innervating NDNF-cells are in SP (with typical pyramidal cell morphology) and in SO (likely O-LM interneurons) ^31,32^. NDNF-cells may also be modulated by neurons at the medial septum and locus coeruleus, which house cholinergic, noradrenergic and dopaminergic neurons. This widespread connectivity suggests that NDNF-cells may integrate information from various regions to SLM, as suggested by recent electrophysiological analysis ^44^.

Regarding outputs, besides directly inhibiting the distal dendrites of CA1 pyramidal cells, NDNF-cells inhibit the GABAergic neurons in SR (Fig. 2 and Fig. 7). NDNF-cells may also innervate GABAergic neurons in the pyramidal layer and SO as well, since their axonal terminals are found there. This dis-inhibits the CA3 input to CA1 at SR and pyramidal layer, which leads to multiple consequences in local computation, such as strengthening the CA3-to-CA1 inputs, facilitating the induction of synaptic plasticity of CA3-to-CA1 synapses ^32^, or contributing to oscillations ^56,57^. Together, the data indicate that NDNF-cells coordinate the dual inputs to CA1 pyramidal cells.

### 2. SLM neurogliaform cells coordinate memory encoding and retrieval

How is learning and memory regulated by NDNF-cells mediated inhibition/dis-inhibition of the dual inputs? In hippocampal circuits, CA1 is where memory encoding and retrieval converge^6,16–18^. Plasticity at CA3-to-CA1 synapses is predicted to be essential for memory encoding and storage ^58–60^. Indeed, learning triggers plasticity in these synapses^61^, and silencing CA3-to-CA1 inputs impairs memory ^62^. On the other hand, the EC3 inputs to CA1 are generally believed to convey online sensory information to CA1 and can serve as cues for memory retrieval ^59,63^. The above model by Hasselmo and colleagues is consistent with this functional division between these two major inputs. However, it is not clear why CA3-to-CA1 synapses could have strong synaptic potentiation for memory encoding when the transmission of these synapse is weak. After all, the lack of “cooperativity” among synapses when synaptic transmission is weak should be detrimental to LTP induction ^28^. Furthermore, when CA1 neuronal spikes temporally align with EC3 inputs, CA3-to-CA1 synaptic activity occurs 20-30 ms after the spikes. This may inhibit synaptic potentiation according to the spike-timing-dependent plasticity (STDP) rules^64^.

The weak transmission and the simultaneous need for synaptic plasticity can be reconciled by NDNF-cells-mediated inhibition of EC3- and disinhibition of CA3-inputs. During encoding, high activity of EC3 inputs will activate NDNF-cells (Fig.9). The activation of NDNF-cells dis-inhibits the CA3-to-CA1 pathway. Disinhibition is a powerful mechanism for strengthening synaptic potentiation at CA3-to-CA1 synapses as well as at synapses in many other circuits ^32,51^. Dis-inhibition also increases the possibility for CA3 inputs to fire CA1 pyramidal neurons and therefore increase the possibility of synaptic potentiation through STDP mechanism. Increasing the firing of NDNF-cells will then facilitate the plasticity at CA3-to-CA1 synapses and memory encoding. On the other hand, increased NDNF-cell activity strongly inhibits EC3 inputs and impairs their function as cues to retrieve memories. Therefore, the NDNF-cells offer a bridge between the two major inputs to CA1 and work critically in both encoding and retrieval. Their activity level sets CA1 in either encoding or retrieval mode to minimize the interference between the two processes and to protect memory-related synaptic plasticity from unwanted alterations.

**Figure 9.**
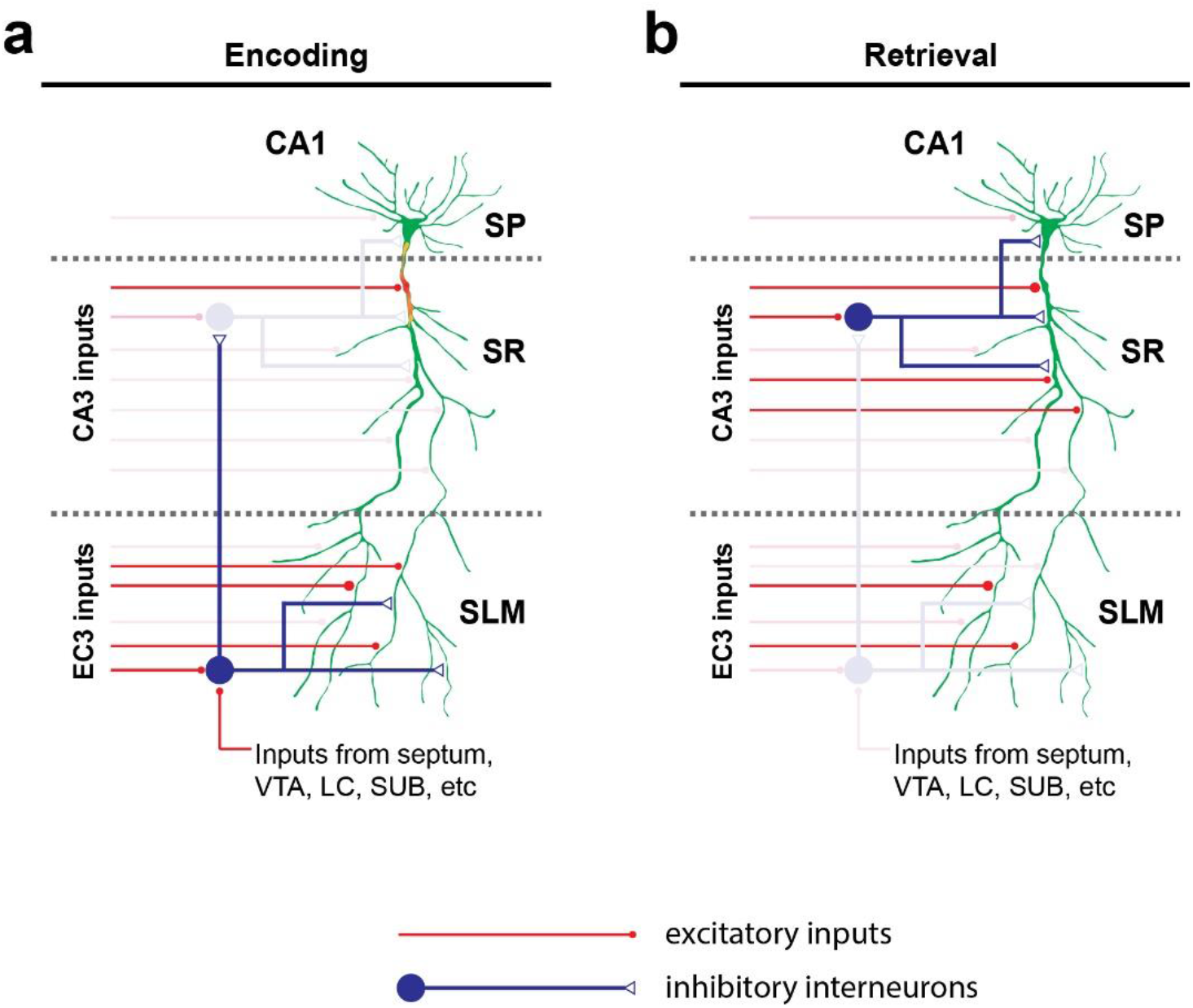
NDNF-cells coordinate encoding and retrieval in hippocampal local circuits. CA1 pyramidal neurons (green) receive major excitatory inputs (red) from EC3 and CA3. Memory encoding preferentially occurs with strong EC3 inputs, while retrieval occurs with strong CA3 inputs^13^. (**a**) When EC3 inputs are strong, they activate NDNF-cells (blue cell at SLM), which in turn regulate the local circuit by direct inhibition at SLM, and dis-inhibition at SR through inhibiting SR inhibitory interneurons (blue cells). This disinhibition facilitates the induction of synaptic potentiation of CA3-to-CA1 synapses for memory encoding. (**b**) Memory retrieval preferentially occurs in the theta phase when synaptic transmission of CA3 inputs are strong and EC3 inputs are weak. Without the dis-inhibition by NDNF-cells, the CA3 inputs activate feedforward inhibition by recruiting SR interneurons, which enables accurate retrieval of information while simultaneously reducing the possibility of altering synaptic strength.

### 3. Limitations and future studies

Although our results demonstrate that the NDNF-cells play distinct roles in memory encoding and retrieval, there are certainly a few weaknesses in the data and some missing links awaiting further research. Firstly, calcium imaging lacks the temporal resolution to examine NDNF-cell activities and their synaptic actions relative to the phases of theta oscillation. *In vivo* electrophysiological recording of NDNF-neurons with optogenetic tagging may shed more light on this. Secondly, our behavioral experiments were conducted with excitatory DREADD system. Our attempts at loss-of-function analysis of NDNF-cells, including with inhibitory DREADD or optogenetics did not provide a clear conclusion, possibly because these approaches did not inhibit NDNF-cells sufficiently, or non-NDNF-cells at SLM compensated. This problem may be solved by further optimizing the AAV tools or develop approaches to target additional SLM interneurons. Thirdly, although our retrograde tracing showed broad innervation of NDNF-cells, we do not know how these inputs regulate the activity of NDNF-cells. In the future, we can examine the environmental factors and circuit mechanisms that activate NDNF-cells, as well as how the regulation of pyramidal cells by NDNF-cells is coupled with the activity in downstream brain circuits. In addition, although previous work has demonstrated that the disinhibition at SR could effectively facilitate LTP there^32^, recording of LTP while activating/inhibiting NDNF-cells may provide more direct evidence.

In summary, by selectively targeting and controlling the NDNF-positive interneurons at SLM, we identified their physiological features and crucial contributions to learning and memory in behaving animals. Further exploration of these cells will elucidate the cognitive processing algorithms behind memory encoding and retrieval.

## Acknowledgments

This study was supported by Klingenstein-Simons Fellowship Awards in Neuroscience (to WX) and grant from NIH/NINDS (NS104828 to WX). We thank Dr. Hongkui Zeng (Allen brain institute) for providing the Ndnf-IRES2-dgCre mouse line, Ms. Wenqin Rita Du and Ms. Elizabeth Li for preparing AAVs. We thank the Metabolic Phenotyping Core (Touchstone Diabetes Center) and Neuroanatomy/Histology/Brain Injection Core (Center for Hypothalamic Research) (supported by NIH funding P01-DK119130-02 to Laurent Gautron) for RNAscope analysis. We also thank Dr. Denise Ramirez, Dr. Julian Meeks and the Whole Brain Microscopy Facility at UT Southwestern; and Dr. Shin Yamazaki (Neuroscience Microscopy Facility at UT Southwestern) for help with imaging.

## Author contributions

WX supervised the research; JG, HCO, SJO and WX designed the research and conducted the experiments; all authors participated in analyzing the results and writing the paper.

## Materials and Methods

### Mice

Heterozygous Ndnf-IRES2-dgCre-D knock-in mice (JAX stock No. 028536) were bred with C57BL/6J mice (UT Southwestern breeding core) to generate heterozygous Cre+ mice and Cre− littermates. Some Ndnf-Cre + mice were crossed with Ai9 mice (JAX stock No. 007909) to visualize Cre expression. Mice were housed together (5 mice per cage) on a 12 hr light /12 hr dark cycle with ad libitum access to food and water. Experiments were carried out during the light cycle. Male and female mice were used for histology, rabies virus tracing and electrophysiology experiments. Male mice were used for behavior experiments. All animal procedures were in accordance with protocols approved by UT Southwestern Institutional Animal Care and Use Committee and complied with the Guide for the Care and Use of Laboratory Animals by the National Research Council.

### Viral vector construction and preparation

AAV vectors were constructed with AAV2 inverted terminal repeats (ITRs). In AAV hDlx-DIO-EGFP, the following components were arranged sequentially downstream of left-ITR: hDlx enhancer and a minimum promoter, loxp site and lox2272 site, inverted open reading frame of EGFP, Loxp site and Lox2272 site followed by hGH poly A sequence, and the right ITR. The AAV hDlx-DIO-TVA-EGFP, AAV hDlx-DIO-hM3Dq-mCherry, AAV hDlx-DIO-ChIEF-EGFP was constructed by replacing the open reading frame of EGFP in AAV hDlx-DIO-EGFP with TVA-EGFP, hM3Dq-mCherry, or ChIEF-EGFP, respectively.

AAVs were packaged with AAV-DJ capsids except otherwise stated. Virus was prepared with iodixanol gradient centrifugation. Briefly, AAV vectors were co-transfected with pHelper and pRC-DJ into AAV-293 cells. Cells were collected 72 hr later, lysed, and loaded onto iodixanol gradient for centrifugation at 400,000 g for 2 hr. The fraction with 40% iodixanol was collected, washed, and concentrated with 100,000 MWCO tube filter. The genomic titer of virus was measured with quantitative real-time PCR. The titers of AAVs used for stereotaxic injection were in the range of 0.5-2 × 10^13^ copies/ml.

### RNAScope multiplex fluorescent in situ hybridization

Mouse brains were fixed through intracardiac perfusion with 10% formalin. Their brains were post-fixed 24 hours at 4°C. After overnight 30% sucrose incubation, 25-μm coronal sections were prepared with a freezing microtome. The tissue was pretreated with H2O2, mounted on SuperFrost slides, and desiccated for approximately one hour. On the following day, the tissue was processed for ISH following the manufacturer’s instructions for multiplex ISH (Advanced Cell Diagnostic; cat# 323100). The tissue was pretreated with a protease-based solution followed by hybridization with the appropriate probes at 40°C for 2 hours. The RNAscope probes used were cat# 447471-C2 (Mm-Ndnf-C2), cat# 437651-C3 (Mm-Nos1-C3), cat# 400951 (Mm-Gad1), cat# 402271 (Mm-cck). Signal amplification was achieved using Opals 520 and 620 diluted at 1/1,500. Lastly, fluorescently-labeled brain sections were counterstained with DAPI, coverslipped with Ecomount, and stored at 4°C.

### Stereotactic surgery

NDNF-Cre+ and Cre− littermates (6-10 weeks) were used for surgeries. All surgeries took place in aseptic conditions while mice were anaesthetized with tribromoethanol (125–250 mg/kg) in a manual stereotactic frame (Kopf). Mice were allowed to recover for 14 - 28 days and single housed before behavioral testing.

To visualize NDNF-expressing interneurons in CA1, we injected 500 nL of AAV hDdlx-DIO-EGFP. The viral solution was injected bilaterally into the dorsal CA1 of NDNF-Cre+ mice in stereotactic surgery as described above, at coordinates −1.95 anteroposterior (AP), +/-1.25 mediolateral (ML), and 1.45 dorsoventral (DV), in millimeter measured from bregma on the skull surface. Injections were performed using a 5 ml Hamilton syringe through a hydraulic pump (Harvard Instruments). Injection speed was 100 nl/min, and the injection needle was slowly raised 5 minutes after completion of virus delivery.

For monosynaptic retrograde neuronal tracing, a total volume of 500 nl of two helper viruses mixed by 2:1 (AAV hDlx-DIO-TVA-EGFP and AAV CAG-DIO-oG) were unilaterally injected into the dorsal CA1 of NDNF-Cre+ mice at coordinates AP: −1.95, ML: +1.25 and DV: 1.45. Three weeks later, 500 nl of a RV*dG* pseudotyped with the envelope protein from avian ASLV type A (EnvA) and fused with mCherry was injected to the same region (Salk Viral Vector Core).

For electrophysiology experiments to confirm EC inputs of NDNF cells, we injected 500 nl of a AAV hDlx-DIO-EGFP into dorsal CA1 to visualize NDNF-cells and AAV hSyn-DIO-ChIFF and AAV hSyn-Cre mixed with 3:1 ratio at bilateral EC (AP:-4.6, ML: +/−3.5, and DV: 2.6). For electrophysiologically confirming SUB inputs of NDNF cells, we injected 500 nl of a AAV hDlx-DIO-EGFP into dorsal CA1 to visualize NDNF-cells and AAV hSyn-DIO-ChIEF and Lentivirus FUW-Cre mixed with 2:1 ratio at bilateral SUB (AP:−3.08, ML: +/−1.4, and DV: 1.65).

For cell-type-specific chemogenetic excitation of NDNF-expressing interneurons, we bilaterally injected 500 nl of AAV hDlx-DIO-hM3Dq-mCherry in dorsal CA1 of NDNF-Cre+ and Cre− littermates. Optogenetic excitation for electrophysiology recordings required bilateral injections of AAV dlx-DIO-ChIEF-EGFP in dorsal CA1 of NDNF-Cre+ mice. For global activation of CA1 interneurons, C57BL/6J mice were assigned randomly to control and experimental group. AAV hDlx-DIO-hM3Dq-mCherry and AAV hSyn-Cre were mixed and 750 nl of virus was injected into bilateral dorsal CA1 region into experimental group. 750 nl of AAV hDlx-DIO-hM3Dq-mCherry was injected into control group.

### Histology and microscopy

Mice were transcardially perfused with 10 ml of PBS followed by 40 mL of 4% paraformaldehyde (PFA) in PBS. The brains were extracted and postfixed overnight in 4% PFA at 4°C, and cryoprotected in 30% sucrose. Brains were sectioned with a cryostat to a thickness of 40 μm. Free-floating sections were washed in PBS, incubated with DAPI (1 μg/mL) and mounted on slides. The whole-mount brain sections were scanned with Zeiss AxioscanZ1 digital slide scanner with a 10X objective. The high-resolution images were taken with ZEISS LSM 880 with Airyscan confocal microscope.

### Immunohistochemistry

After perfusion and sectioning, brain slices were washed three times using PBS, permeabilized and then incubated with blocking solution (5% normal goat serum in 0.1% Triton X-100 in PBS) for 1 hr followed by primary antibody incubation overnight at 4°C using goat Anti-nNOS (ab1376, Abcam, 1:300) antibodies. The next day, slices were washed with PBS and incubated with appropriate secondary antibodies for 2 hr (1:500, donkey anti-goat Alexa Fluor 633).

### Pharmacogenetic manipulation

Clozapine (0444, Tocris Bioscience) was first dissolved in dimethylsulfoxide (DMSO, D2438, Sigma, final concentration of 32.6mg/ml). Aliquots were stored at −20°C and equilibrated to room temperature before being further diluted in 0.9% sterile saline solution to a final concentration of 0.01 mg/ml.

Saline (0.9% NaCl) or clozapine dissolved in saline (0.1 mg/kg body weight) was injected intraperitoneally into mice 30 min before behavioral experiments. After injection, mice were left undisturbed in their home cage until the start of the experiment.

### Contextual fear conditioning

Mice were handled in 1-min epochs every day for 5 days before behavioral training. Clozapine or saline was intraperitoneally injected 30 min before training, right after training or 30 min before recall test according to the purpose of the experiment. During training, each mouse was allowed to explore the chamber without any stimulation for 120 s. The chamber had metal walls, metallic bars on the floor, and 10% ethanol odorant. A tone amplified to 82-83 dB and lasting 30 s was presented at 120 s after mouse entering context. The last 2 s of the tone were paired with an electric footshock (2 s, 0.8 mA). The mouse was removed from the chamber 30 s after the end of shock. On day 2, 24 hr after the training session, contextual fear memory was assessed by placing the mouse in the original context for 300 s and measuring the percentage of total time that the mouse spent freezing. On day 3, mice were exposed to an altered context (decorated walls, smooth floor surface, and 5% vanilla odorant). Mice were monitored for 300 s for freezing to the modified context. The percentage of time that each mouse spent freezing was determined using FreezeView software (Version 2.26, Actimetrics Inc, Wilmette, IL, USA). For all the behavior experiments, the experimenters were blind to the group assignment during handling and behavior tests.

### Novel place recognition

The paradigm was divided in three phases. First, animals were handled for 5 consecutive days and then habituated to a decorated open field (monochromatic horizontal lines and polychromatic straight lines on opposite walls) for two consecutive days (10 min sessions). Second, during the familiarity phase, animals were exposed after clozapine i.p. Injections to two objects in fixed positions approximately 7 cm away from the decorated walls (10 min sessions). Finally, 5 hrs (when clozapine was injected before test) or 24 hrs (when clozapine was injected before training) later, animals were tested in the same environment, but with one object placed in a different position (10 min sessions). A video tracking system (ANY-maze, Stoelting) was used to track the head-position of animals and analyze the time spent exploring the circumference of each object (around 5 cm diameter). The arena and objects were cleaned with water before each trial to remove odorant cues. If an animal spent less than 20 s exploring objects during either training or test (10-min session each), the data was excluded from further analysis.

### Novel object recognition Test

Animals were handled for 5 consecutive days and then habituated to a decorated open field. To activate NDNF-cells during learning, animals received clozapine i.p. injections and then were exposed to two identical objects in fixed positions approximately 7 cm away from the decorated walls (10 min sessions). Finally, animals were tested in the same environment 24h later (to guarantee clozapine had already been metabolized), but with one object replaced by a new one (10 min sessions). Objects were interchanged to avoid animal preference due to position in the open field. To activate NDNF interneurons during memory recall, animals received i.p. clozapine injections before the test session (5h after training). A video tracking system (ANY-maze, Stoelting) was used to track the head-position of animals and analyze the time spent exploring the circumference of each object (around 5 cm diameter). The arena and objects were cleaned with water before each trial to remove odorant cues. If an animal spent less than 20 s exploring objects during either training or test (10-min session each), the data was excluded from further analysis.

### Open field

Thirty minutes after clozapine injection, locomotor activity was evaluated by placing one mouse at a time into an opaque plastic (50×50 cm) open-field arena and allowing the mouse to explore for 10 min. Activity in the open-field was recorded and quantified using the ANY-maze video tracking system.

### Elevated plus maze

Thirty minutes after clozapine injection, a mouse was put in the center of the maze to freely explore for 10 minutes. The maze was made of beige plastic consisting of two opposite open arms (66 × 5.1 cm) and two opposite closed arms surrounded by walls. The center section between arms consisted of a square with dimensions of 5.1 × 5.1 cm. The maze was elevated 38.7 cm above ground. The trials were recorded in video and analyzed with ANY-maze software. All behavioral instruments were acquired from San Diego Instruments, Inc.

### Brain slice electrophysiology

3-4 weeks after AAV injection, transverse or coronal slices of the dorsal hippocampus (300 μm) were prepared with a vibratome (Leica VT1200) in ice cold cutting solution containing (in mM): 2.5 KCl, 1.2 NaH_2_PO_4_, 26 NaHCO_3_, 10 D-glucose, 213 sucrose, 5 MgCl_2_, 0.5 CaCl_2_. The slices were incubated for 30 min in artificial cerebrospinal fluid (ACSF) containing (in mM): 124 NaCl, 5 KCl, 1.2 NaH_2_PO_4_, 26 NaHCO_3_, 10 D-glucose, 1.3 MgCl_2_, 2.5 CaCl_2_ at 32 °C and then for at least 1 h at room temperature. The cutting solution and ACSF were adjusted to pH 7.3-7.4 and 290 - 300 mOsm and constantly aerated with 95% O_2_/5% CO_2_. Whole-cell patch clamp recording was performed in a recording chamber perfused (∼1 ml/min) with oxygenated ACSF at 26-28 °C. For most of the intracellular recordings, the recording pipettes (2.5-4 MΩ) were filled with internal solution containing (in mM): 125 K-gluconate, 20 KCl, 4 Mg-ATP, 0.3 Na-GTP, 10 Na_2_-phosphocreatine, 0.5 EGTA, 10 HEPES, adjusted to pH 7.3-7.4 and 310 mOsm. For the recordings shown in Fig. 6F, the recording pipettes (2.5-4 MΩ) were filled with internal solution containing (in mM): 138 K-gluconate, 7 KCl, 4 Mg-ATP, 0.3 Na-GTP, 10 Na_2_-phosphocreatine, 0.5 EGTA, 10 HEPES, adjusted to pH 7.3-7.4 and 310 mOsm. For optogenetic experiments, blue light (473 nm) was delivered by LED coupled with a 40x water objective. For electrically stimulating SR or SLM fibers, a focal stimulating pipette (∼ 1 MΩ) was filled with ACSF and placed at 50 μM below the surface of the slice. For fIPSP or fEPSP recording, a pipette (∼ 1.5 MΩ) was placed at different sublayers of CA1.

To measure early or late spiking, current step was first adjusted to below firing threshold and then increased by every 5 pA until one action potential was triggered. To produce persistent firing, 20 1-s current step at 200 pA with 1-s interval was delivered into recorded cell. If the cell showed spontaneous action potentials during the interval, the current step protocol was terminated and spontaneous action potentials were recorded without any stimulation.

### In vivo calcium imaging

In preparation for calcium imaging in NDNF-interneurons, NDNF-Cre+ mice were unilaterally injected with 500 nl of AAV9 hSyn-FLEX-jGCaMP7f in dorsal CA1. Three weeks after injection, a GRIN lens of 0.5 mm in diameter was implanted on top of SLM of CA1 at AP: −1.95, ML: 1.25, DV: 1.45. The lens was protected by a plastic tube. At least 2 weeks after lens implantation, the plastic tube was removed and a metal base plate was attached to the skull. A custom-built mini-endoscope was installed and used for calcium imaging ^65^. Mice were tested with contextual fear conditioning task as described above.

For calcium events recorded in CA1 pyramidal cells during the excitation of NDNF-interneurons, animals were unilaterally injected with 500 nl of AAV hSyn-GCaMP6s-P2A-NLS-dTomato. Additionally, these mice received bilateral delivery of AAV hDlx-DIO-hM3Dq-mCherry in dorsal CA1 as described above. A GRIN lens was implanted after viral injection at AP: −1.95, ML: 1.25, DV: 1.25. The same procedure was performed as described above. For imaging CA1 pyramidal layer, mice were imaged in home cage and a 1-min session was recorded at −20, −10, −1, 10, 20, 30, 40, 50, 60, 70, 80, 90, 120, 180 min before (-) or after saline or 0.1 mg/kg clozapine injection. At least a week after home cage imaging, mice were injected with 0.1 mg/kg clozapine and trained for contextual fear conditioning as described above. Saline was injected before context and altered context tests.

For calcium imaging of CA1 presynaptic inputs from EC or CA3, C57BL/6J mice were unilaterally injected with 500 nl of AAV-hSyn-Syb2-GCaMP6f and highly diluted AAV-hSyn-tdTomato (50:1) at AP: −4.8, ML:3.5, DV: 2.6 (EC) or bilaterally at AP: −2.35, ML: +/−2.6, DV:2 (CA3). A GRIN lens was implanted at AP: −1.95, ML:1.35, DV: 1.5 on the top of SLM layer for EC input imaging or at AP: −1.95, ML:1.25, DV: 1.3 on the top of SR layer for CA3 input imaging. The procedure of endoscope installation was the same as described above.

### Calcium imaging data analysis

For NDNF neurons imaging, the rigid motion was corrected by NoRMCorre ^66^. The frame by frame large-scale background was estimated by applying a 20-pixel gaussian filter in ImageJ. The ROI for individual neuron was manually selected in ImageJ and the location background of each ROI was defined as the average intensity in an annulus whose inner diameter was the diameter of ROI and outer diameter is around 2-fold of the diameter of ROI. The large-scale and local backgrounds were subtracted from the average intensity within ROI to obtain the signal of individual ROI. The fluorescence background (F) for each session was defined by the average of the lowest 20% of signal value within a given session. After obtaining dF/F value, signal was normalized to the maximum value for further quantification.

For CA1 pyramidal cells imaging, the rigid motion was corrected by NoRMCorre and the calcium signal and deconvoluted events were extracted and analyzed with the extended CNMF for microendoscopic data (CNMF-E) approach ^67^. For the imaging in home cage in Fig. 5C to E. Calcium event rate during each recording session was calculated by dividing the number of events within this particular session by the number of all neurons detected in all session. The calcium event rate for each session was further normalized to the average of first 3 sessions (before injection). For the imaging during fear conditioning, in Fig. 4G calcium event rate for individual neuron was pooled with a 2-s bin. In Fig. 4H, the averaged neuronal activity of first two minutes in training was defined as exploration time.

For calcium imaging of CA1 presynaptic inputs, raw imaging data was loaded into ImageJ. The calcium sign was measured in a ROI covering ∼80% of lens area. The moving background (F) for each frame was measured by the minimum signal intensity within 10-s before this frame. The minimum signal intensity of the first 10 s of recording was used as the background of the frames in the first 10 s. To compare the signal between EC input and CA3 input after shock, the dF/F value was first normalized the maximum dF/F value over the whole recording. Then, the area under the curve (AUC) was measured with a 5-s bin. The binned AUC was further normalized to the average value of first 24 bins, corresponding to first 120 s in training.

### Quantification and Statistical Analysis

Data are presented as mean ± standard error of the mean (SEM). Sample number (n) indicates the number of cells, slices or mice in each experiment and is specified in the results or the figures. The sample number (n) used for statistics analysis is from biological replications. The methods for statistical analysis are described in the results or the figures. No explicit power analysis was used to determine the sample size. The sample size was decided during experimental design based on previous literatures on similar experiments.

## Supplemental Information

### The supplemental information includes

Figs. S1 to S8

### Other supplementary materials for this manuscript include

Movies S1-S3

Movie S1: Representative video showing the activity of NDNF-cells. The video is at 5X speed.
Movie S2: Representative video showing the neuronal activity at CA1 pyramidal layer. The video is at 5X speed.
Movie S3: Representative video showing the calcium signal of bulk EC3-CA1 synaptic terminals. The video is at 4X speed.

## Supplementary figures

**Fig. S1:**
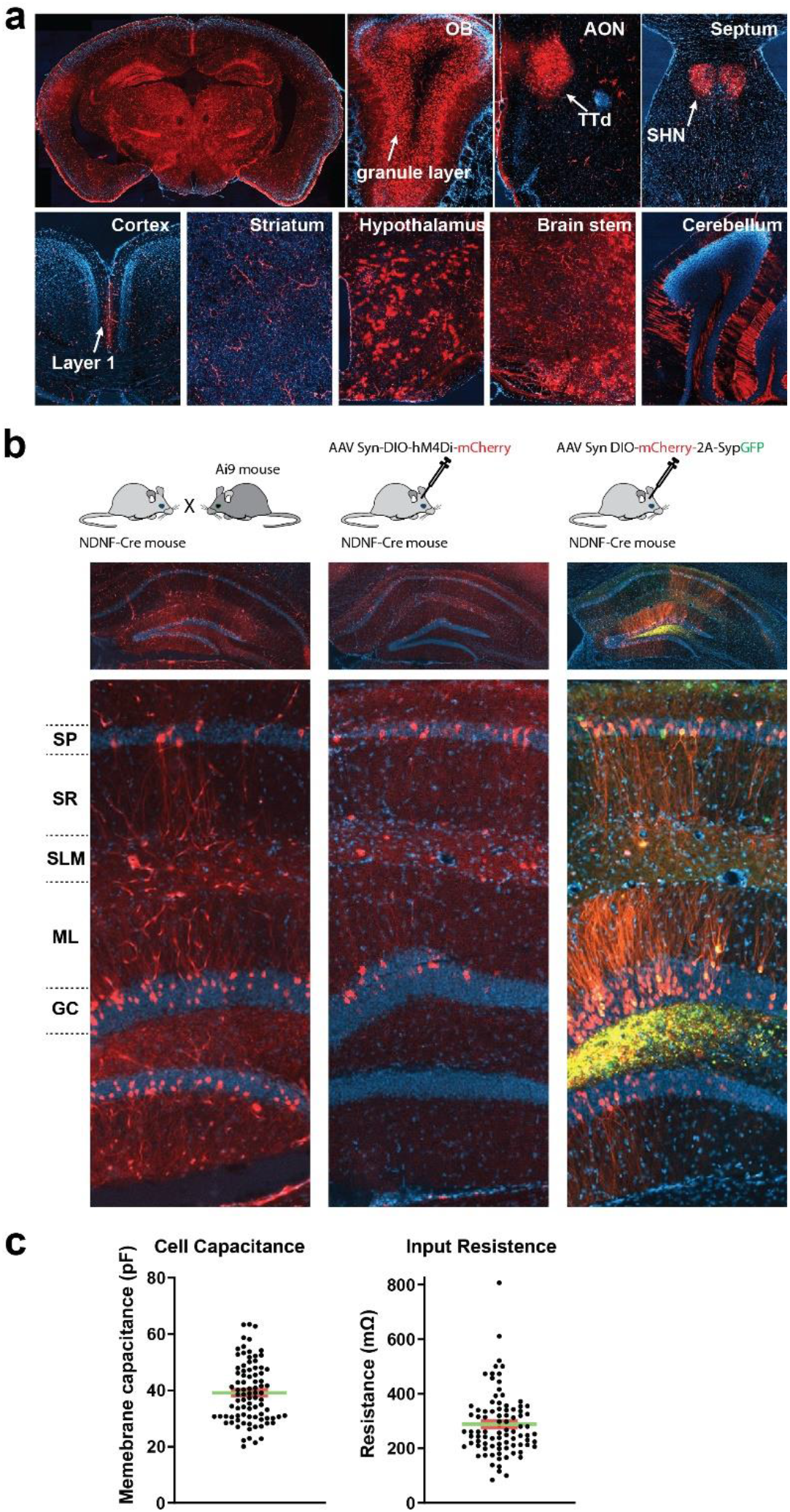
Brain-wide expression pattern of NDNF-Cre. (**a**) Brain sections from a NDNF-Cre mouse crossed with a tdTomato report mouse (Ai9). The brain was fixed at the age of 2 months. The representative coronal sections showed the widespread expression of Cre in the brain, including in granule layer of olfactory bulb, Taenia tecta (TTd), septo-hippocampal nucleus (SHN) in septum, layer1 interneurons in cortex, striatum, glia cells in hypothalamus, glia cells in brain stem and Purkinje cells in the cerebellum. Blood vessels are labeled in all the regions examined. (**b**) NDNF-Cre mice were either crossed with Ai9 report mouse (left panel) or injected with AAV mediating Cre-dependent expression of hM4Di-mCherry into the hippocampus (middle panel), or injected with AAV mediating the expression of mCherry and EGFP fused to synaptophysin into the hippocampus (right panel) (Top row). Representative coronal sections of the hippocampus (middle row), and the enlargement of the CA1 region (bottom row) were shown. Besides the interneurons localized at SLM, the pyramidal cells at the CA1 and the granule cells at the DG also expressed Cre. (**c**) Passive membrane properties (cell capacitance and input resistance) of NDNF-cells.

**Fig. S2:**
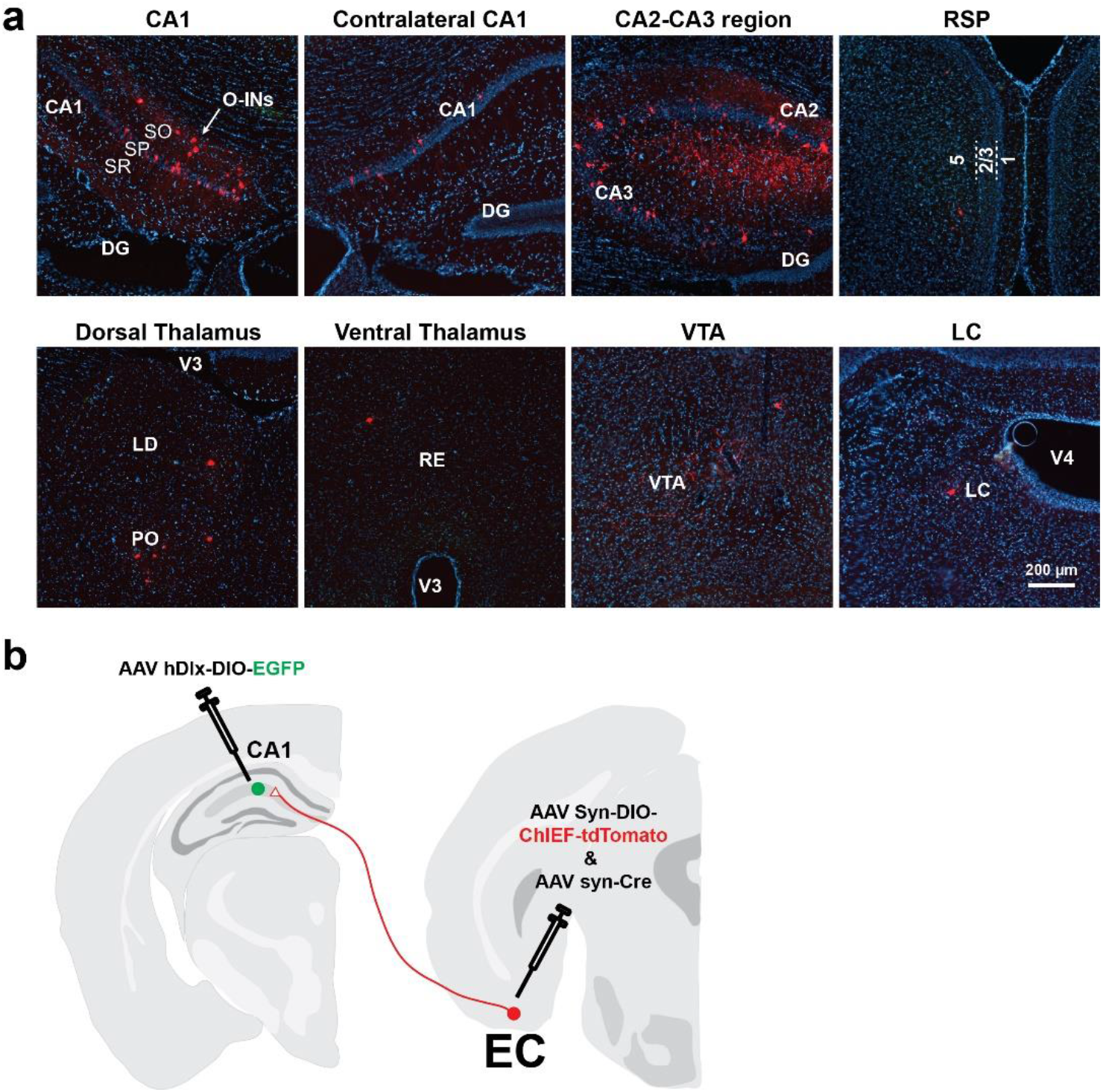
Synaptic innervation of NDNF-cells. (**a**) Additional photos showing the presynaptic input regions/cells of NDNF-cells traced with recombinant rabies virus. (**b**) Experimental design for expressing ChIEF in EC-CA1 synapses and expressing EGFP in NDNF-cells.

**Fig. S3:**
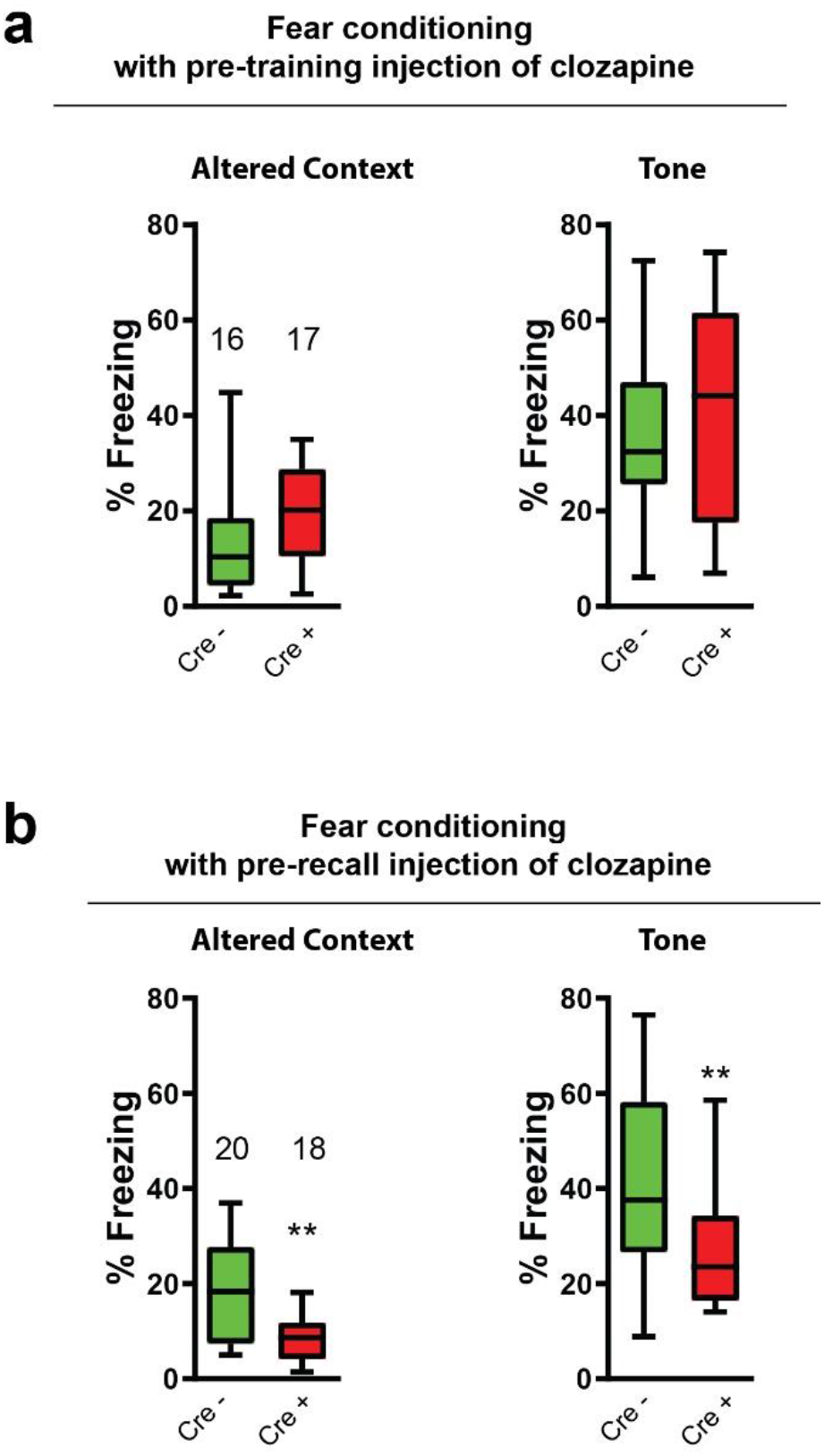
Freezing to altered context and tone. (**a**) Freezing to altered context or tone of the mice shown in Fig.4c-d. Clozapine was not administrated before the tests. (**b**) Freezing to altered context or tone of the mice shown in Fig.4i-j. Clozapine was administered before the tests.

**Fig. S4:**
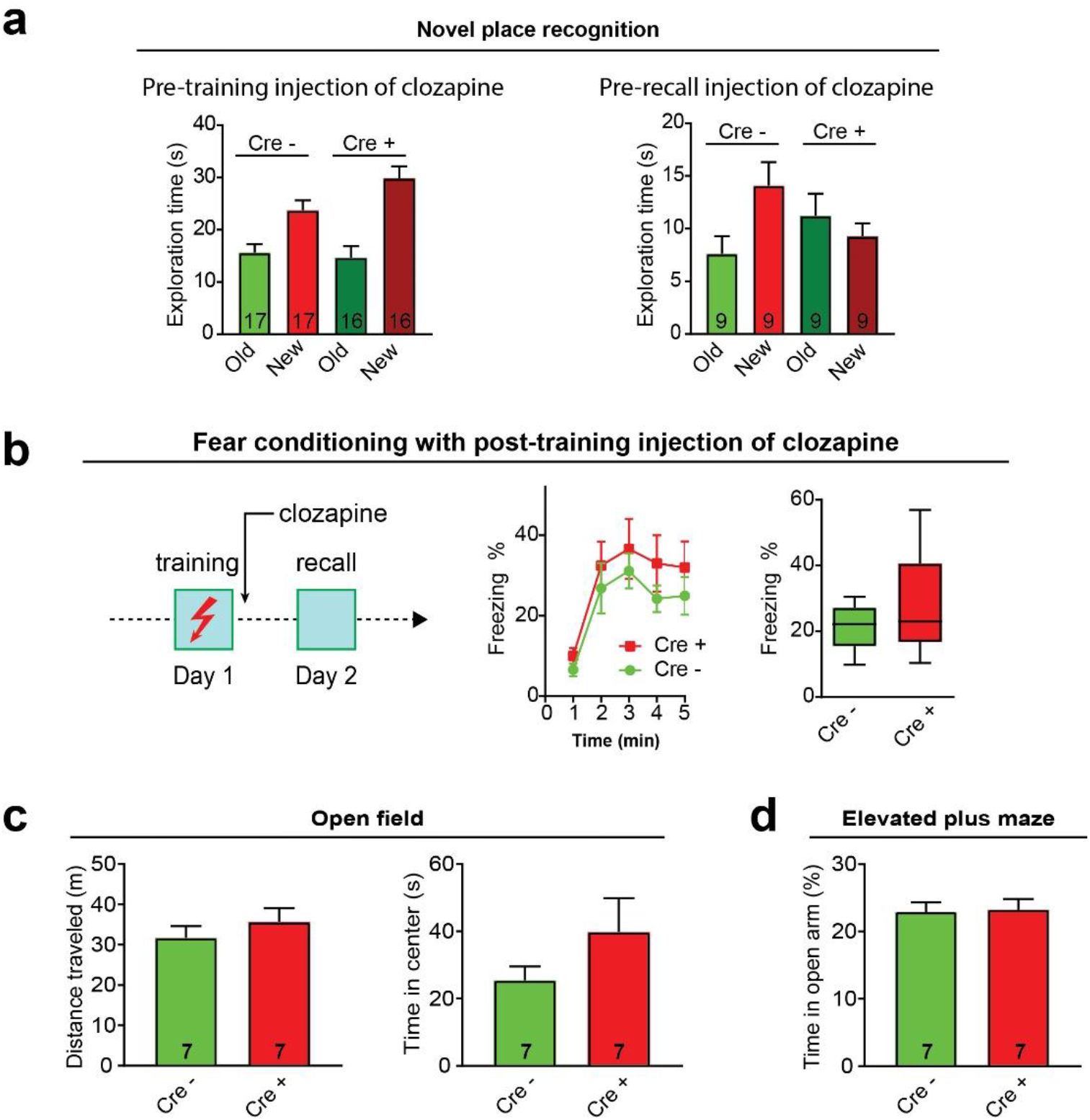
Additional behavioral tests with NDNF-cell activation. (**a**) Exploration time for new and old locations in novel place recognition test. The left panel corresponds to data shown in Fig. 4h. The right panel corresponds to data shown in Fig. 4k. (**b**) Contextual fear conditioning with clozapine been injected right after training. (**c**) Left: Distance traveled; right: Time in the center in open field test when clozapine was injected 30 min before test (p=0.41, student t-test for distance traveled; p=0.21, student t-test for time in center). (**d**) Percentage of time in the open arm in elevated plus maze test when clozapine was injected 30 min before test (p=0.88, student t-test).

**Fig. S5:**
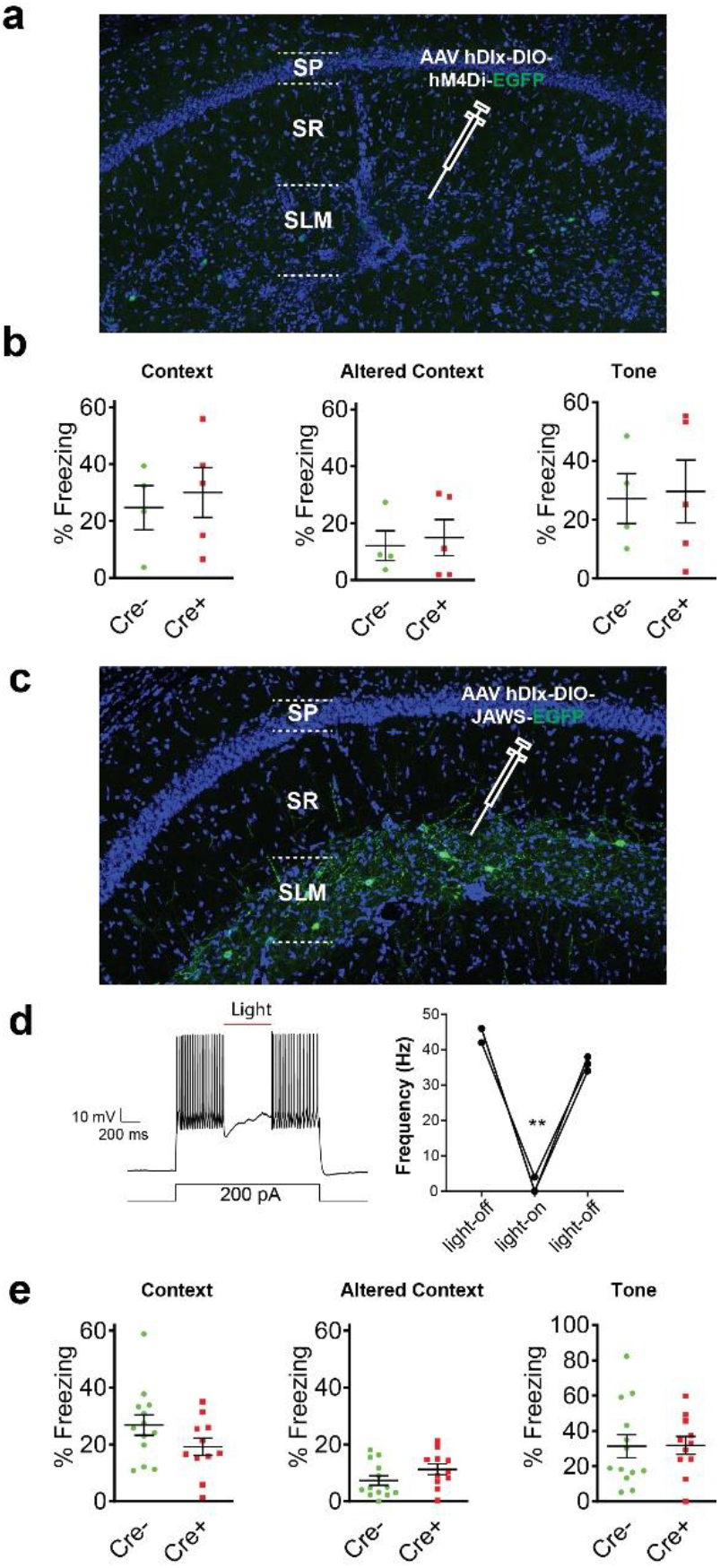
Loss-of-function studies of NDNF-cells. (**a**) Expression of hM4Di-EGFP in NDNF-cells. (**b**) Contextual fear conditioning with clozapine been injected 30min before training. (**c**) Expression of JAWS-EGFP in NDNF-cells. (**d**) Red light inhibited the firing of NDNF-cells. Action potentials were triggered in NDNF-cells by current injections in whole-cell patch-clamp recording mode. (**e**) Contextual fear conditioning with optogenetic silencing of NDNF-cells. Cre− and Cre+ mice were injected with AAV hDlx-DIO-JAWS-EGFP, and implanted with optic fibers in bilateral CA1. Light was delivered during fear conditioning training but not during the tests.

**Fig. S6:**
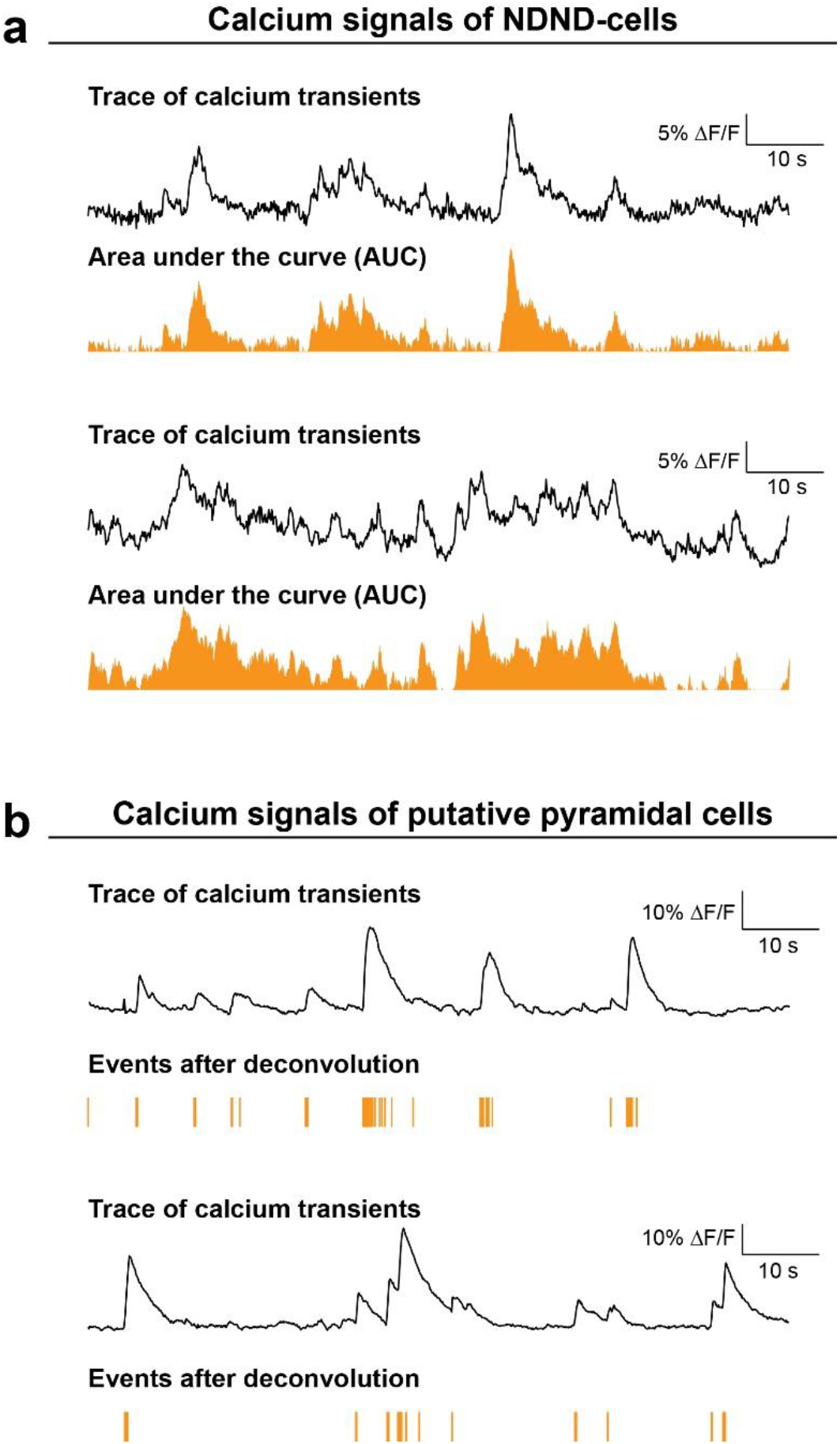
Quantification of calcium signals of NDNF-cells and putative CA1 pyramidal cells. (**a**) Representative traces of NDNF-cells showing the heterogeneous kinetics of calcium signals of NDNF-cells. Due to the difficulty of reducing the calcium signals to individual spikes, we measured the area under the calcium traces (Area under the curve) for quantification. (**b**) Representative traces of calcium imaging of neurons at pyramidal layer—the presumptive pyramidal cells. The calcium transients were deconvoluted into individual firing events with the extended CNMF for microendoscopic data (CNMF-E) method^67^.

**Fig. S7:**
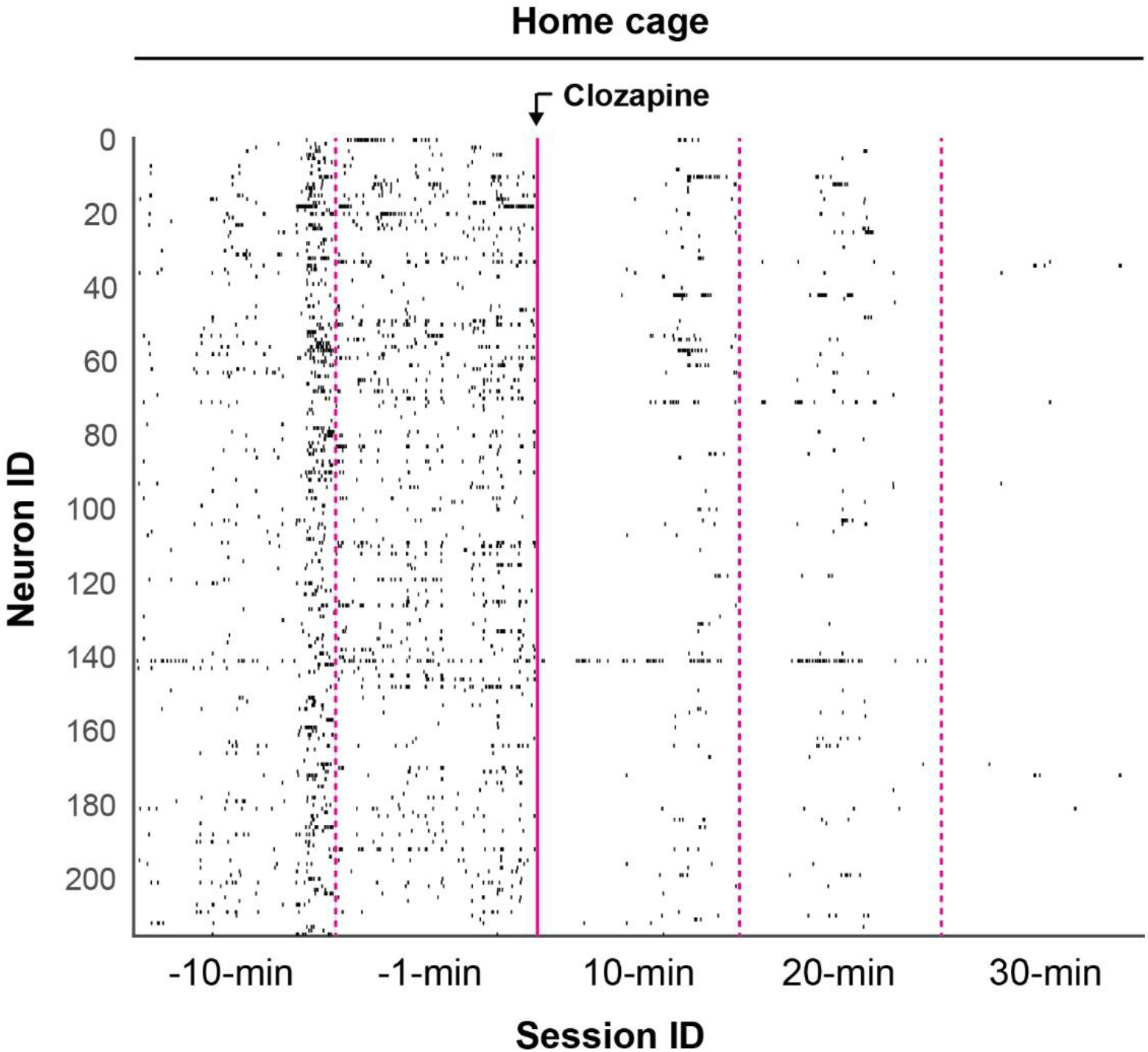
Activation of pan-interneurons in CA1 with hM3Dq inhibits CA1 pyramidal cells.

**Fig. S8:**
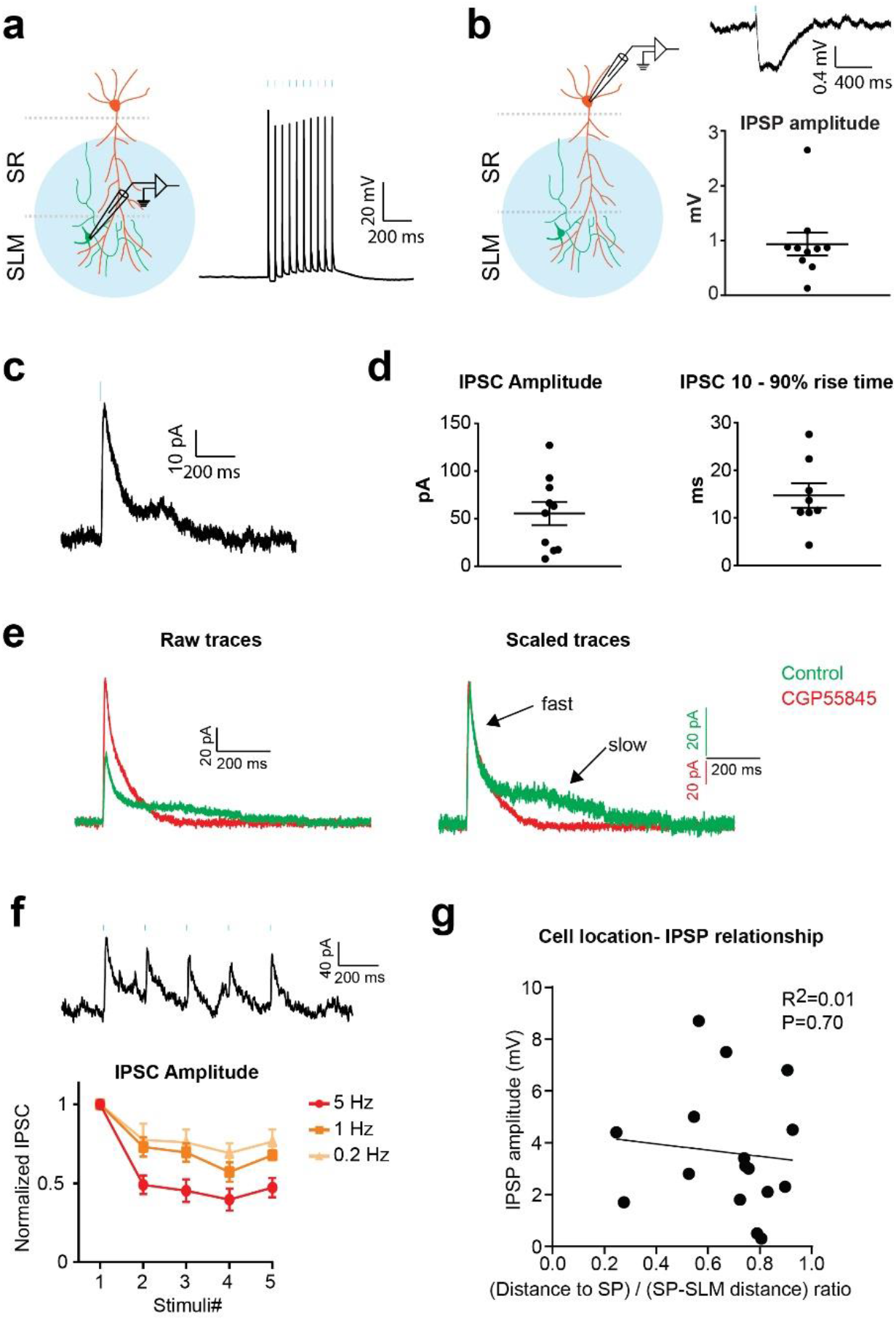
NDNF-Cells inhibition of CA1 pyramidal cells and SR interneurons. (**a**) 2-ms light pulses delivered at 20 Hz reliably triggered action potentials in an EGFP-positive NDNF-Cell. (**b**-**e**) Optostimulation of NDNF-Cells evoked biphasic IPSPs (**b**) or IPSCs (**c**-**e**) in CA1 pyramidal cells. (**e**) GABA_B_ receptors antagonist, CGP55845, blocked the slow phase and increased the amplitude of the fast phase of the IPSCs triggered by photo-stimulation of NDNF-cells. (**f**) Representative trace and quantification of IPSCs recorded in CA1 pyramidal cells evoked by five 2-ms light pulses delivered at 5, 1 or 0.2 Hz to NDNF-cells. The pyramidal cell was held at 0 mV in voltage clamp. The IPSC amplitude was normalized to the amplitude of the first IPSC. (**g**) The amplitudes of the IPSCs recorded from SR interneurons triggered by photo-stimulation of NDNF-cells were not correlated to the locations of the interneurons in SR.

## References

1 Tulving, E. & Thomson, D. M. Encoding Specificity and Retrieval Processes in Episodic Memory. Psychol Rev 80, 352–373, doi:Doi10.1037/H0020071 (1973).

2 Ben-Yakov, A., Dudai, Y. & Mayford, M. R. Memory Retrieval in Mice and Men. Cold Spring Harbor perspectives in biology 7, doi:10.1101/cshperspect.a021790 (2015).

3 Tonegawa, S., Pignatelli, M., Roy, D. S. & Ryan, T. J. Memory engram storage and retrieval. Current opinion in neurobiology 35, 101–109, doi:10.1016/j.conb.2015.07.009 (2015).

4 Frankland, P. W., Josselyn, S. A. & Kohler, S. The neurobiological foundation of memory retrieval. Nature neuroscience 22, 1576–1585, doi:10.1038/s41593-019-0493-1 (2019).

5 Reijmers, L. G., Perkins, B. L., Matsuo, N. & Mayford, M. Localization of a stable neural correlate of associative memory. Science 317, 1230–1233, doi:10.1126/science.1143839 (2007).

6 Tanaka, K. Z. et al. The hippocampal engram maps experience but not place. Science 361, 392–397, doi:10.1126/science.aat5397 (2018).

7 Garner, A. R. et al. Generation of a synthetic memory trace. Science 335, 1513–1516, doi:10.1126/science.1214985 (2012).

8 Liu, X. et al. Optogenetic stimulation of a hippocampal engram activates fear memory recall. Nature 484, 381–385, doi:10.1038/nature11028 (2012).

9 Denny, C. A. et al. Hippocampal memory traces are differentially modulated by experience, time, and adult neurogenesis. Neuron 83, 189–201, doi:10.1016/j.neuron.2014.05.018 (2014).

10 Lisman, J., Cooper, K., Sehgal, M. & Silva, A. J. Memory formation depends on both synapse-specific modifications of synaptic strength and cell-specific increases in excitability. Nature neuroscience 21, 309–314, doi:10.1038/s41593-018-0076-6 (2018).

11 Morris, R. G. et al. Memory reconsolidation: sensitivity of spatial memory to inhibition of protein synthesis in dorsal hippocampus during encoding and retrieval. Neuron 50, 479–489, doi:10.1016/j.neuron.2006.04.012 (2006).

12 Colgin, L. L. et al. Frequency of gamma oscillations routes flow of information in the hippocampus. Nature 462, 353–357, doi:10.1038/nature08573 (2009).

13 Hasselmo, M. E., Bodelon, C. & Wyble, B. P. A proposed function for hippocampal theta rhythm: separate phases of encoding and retrieval enhance reversal of prior learning. Neural computation 14, 793–817, doi:10.1162/089976602317318965 (2002).

14 Tulving, E., Kapur, S., Craik, F. I., Moscovitch, M. & Houle, S. Hemispheric encoding/retrieval asymmetry in episodic memory: positron emission tomography findings. Proceedings of the National Academy of Sciences of the United States of America 91, 2016–2020, doi:10.1073/pnas.91.6.2016 (1994).

15 Roy, D. S. et al. Distinct Neural Circuits for the Formation and Retrieval of Episodic Memories. Cell 170, 1000–1012 e1019, doi:10.1016/j.cell.2017.07.013 (2017).

16 Tayler, K. K., Tanaka, K. Z., Reijmers, L. G. & Wiltgen, B. J. Reactivation of neural ensembles during the retrieval of recent and remote memory. Current biology : CB 23, 99–106, doi:10.1016/j.cub.2012.11.019 (2013).

17 Deng, W., Mayford, M. & Gage, F. H. Selection of distinct populations of dentate granule cells in response to inputs as a mechanism for pattern separation in mice. eLife 2, e00312, doi:10.7554/eLife.00312 (2013).

18 Ji, J. & Maren, S. Differential roles for hippocampal areas CA1 and CA3 in the contextual encoding and retrieval of extinguished fear. Learning & memory 15, 244–251, doi:10.1101/lm.794808 (2008).

19 Hunsaker, M. R. & Kesner, R. P. Dissociations across the dorsal-ventral axis of CA3 and CA1 for encoding and retrieval of contextual and auditory-cued fear. Neurobiology of learning and memory 89, 61–69, doi:10.1016/j.nlm.2007.08.016 (2008).

20 van Strien, N. M., Cappaert, N. L. & Witter, M. P. The anatomy of memory: an interactive overview of the parahippocampal-hippocampal network. Nature reviews. Neuroscience 10, 272–282, doi:10.1038/nrn2614 (2009).

21 Dudman, J. T., Tsay, D. & Siegelbaum, S. A. A role for synaptic inputs at distal dendrites: instructive signals for hippocampal long-term plasticity. Neuron 56, 866–879, doi:10.1016/j.neuron.2007.10.020 (2007).

22 Mizuseki, K., Sirota, A., Pastalkova, E. & Buzsaki, G. Theta oscillations provide temporal windows for local circuit computation in the entorhinal-hippocampal loop. Neuron 64, 267–280, doi:10.1016/j.neuron.2009.08.037 (2009).

23 Douchamps, V., Jeewajee, A., Blundell, P., Burgess, N. & Lever, C. Evidence for encoding versus retrieval scheduling in the hippocampus by theta phase and acetylcholine. The Journal of neuroscience : the official journal of the Society for Neuroscience 33, 8689–8704, doi:10.1523/JNEUROSCI.4483-12.2013 (2013).

24 Fernandez-Ruiz, A. et al. Entorhinal-CA3 Dual-Input Control of Spike Timing in the Hippocampus by Theta-Gamma Coupling. Neuron 93, 1213–1226 e1215, doi:10.1016/j.neuron.2017.02.017 (2017).

25 Schomburg, E. W. et al. Theta phase segregation of input-specific gamma patterns in entorhinal-hippocampal networks. Neuron 84, 470–485, doi:10.1016/j.neuron.2014.08.051 (2014).

26 Siegle, J. H. & Wilson, M. A. Enhancement of encoding and retrieval functions through theta phase-specific manipulation of hippocampus. eLife 3, e03061, doi:10.7554/eLife.03061 (2014).

27 Bieri, K. W., Bobbitt, K. N. & Colgin, L. L. Slow and fast gamma rhythms coordinate different spatial coding modes in hippocampal place cells. Neuron 82, 670–681, doi:10.1016/j.neuron.2014.03.013 (2014).

28 Bliss, T. V. & Collingridge, G. L. A synaptic model of memory: long-term potentiation in the hippocampus. Nature 361, 31–39, doi:10.1038/361031a0 (1993).

29 Pelkey, K. A. et al. Hippocampal GABAergic Inhibitory Interneurons. Physiological reviews 97, 1619–1747, doi:10.1152/physrev.00007.2017 (2017).

30 Royer, S. et al. Control of timing, rate and bursts of hippocampal place cells by dendritic and somatic inhibition. Nature neuroscience 15, 769–775, doi:10.1038/nn.3077 (2012).

31 Lovett-Barron, M. et al. Dendritic inhibition in the hippocampus supports fear learning. Science 343, 857–863, doi:10.1126/science.1247485 (2014).

32 Leao, R. N. et al. OLM interneurons differentially modulate CA3 and entorhinal inputs to hippocampal CA1 neurons. Nature neuroscience 15, 1524–1530, doi:10.1038/nn.3235 (2012).

33 Overstreet-Wadiche, L. & McBain, C. J. Neurogliaform cells in cortical circuits. Nature reviews. Neuroscience 16, 458–468, doi:10.1038/nrn3969 (2015).

34 Klausberger, T. & Somogyi, P. Neuronal diversity and temporal dynamics: the unity of hippocampal circuit operations. Science 321, 53–57, doi:10.1126/science.1149381 (2008).

35 Capogna, M. Neurogliaform cells and other interneurons of stratum lacunosum-moleculare gate entorhinal-hippocampal dialogue. The Journal of physiology 589, 1875–1883, doi:10.1113/jphysiol.2010.201004 (2011).

36 Abs, E. et al. Learning-Related Plasticity in Dendrite-Targeting Layer 1 Interneurons. Neuron 100, 684–699 e686, doi:10.1016/j.neuron.2018.09.001 (2018).

37 Poorthuis, R. B. et al. Rapid Neuromodulation of Layer 1 Interneurons in Human Neocortex. Cell reports 23, 951–958, doi:10.1016/j.celrep.2018.03.111 (2018).

38 Price, C. J. et al. Neurogliaform neurons form a novel inhibitory network in the hippocampal CA1 area. The Journal of neuroscience : the official journal of the Society for Neuroscience 25, 6775–6786, doi:10.1523/JNEUROSCI.1135-05.2005 (2005).

39 Tricoire, L. et al. Common origins of hippocampal Ivy and nitric oxide synthase expressing neurogliaform cells. The Journal of neuroscience : the official journal of the Society for Neuroscience 30, 2165–2176, doi:10.1523/JNEUROSCI.5123-09.2010 (2010).

40 Harris, K. D. et al. Classes and continua of hippocampal CA1 inhibitory neurons revealed by single-cell transcriptomics. PLoS biology 16, e2006387, doi:10.1371/journal.pbio.2006387 (2018).

41 Marion S. Mercier, V. M., Jonathan Cornford, Dimitri M. Kullmann. (bioRxiv 531822; doi: https://doi.org/10.1101/531822, 2019).

42 Dimidschstein, J. et al. A viral strategy for targeting and manipulating interneurons across vertebrate species. Nature neuroscience 19, 1743–1749, doi:10.1038/nn.4430 (2016).

43 Kim, E. J., Jacobs, M. W., Ito-Cole, T. & Callaway, E. M. Improved Monosynaptic Neural Circuit Tracing Using Engineered Rabies Virus Glycoproteins. Cell reports 15, 692–699, doi:10.1016/j.celrep.2016.03.067 (2016).

44 Chittajallu, R. et al. Afferent specific role of NMDA receptors for the circuit integration of hippocampal neurogliaform cells. Nature communications 8, 152, doi:10.1038/s41467-017-00218-y (2017).

45 Varela, C., Kumar, S., Yang, J. Y. & Wilson, M. A. Anatomical substrates for direct interactions between hippocampus, medial prefrontal cortex, and the thalamic nucleus reuniens. Brain Struct Funct 219, 911–929, doi:10.1007/s00429-013-0543-5 (2014).

46 Gomez, J. L. et al. Chemogenetics revealed: DREADD occupancy and activation via converted clozapine. Science 357, 503–507, doi:10.1126/science.aan2475 (2017).

47 Wiltgen, B. J., Sanders, M. J., Anagnostaras, S. G., Sage, J. R. & Fanselow, M. S. Context fear learning in the absence of the hippocampus. The Journal of neuroscience : the official journal of the Society for Neuroscience 26, 5484–5491, doi:10.1523/JNEUROSCI.2685-05.2006 (2006).

48 Chuong, A. S. et al. Noninvasive optical inhibition with a red-shifted microbial rhodopsin. Nature neuroscience 17, 1123–1129, doi:10.1038/nn.3752 (2014).

49 Bezaire, M. J. & Soltesz, I. Quantitative assessment of CA1 local circuits: knowledge base for interneuron-pyramidal cell connectivity. Hippocampus 23, 751–785, doi:10.1002/hipo.22141 (2013).

50 Karayannis, T. et al. Slow GABA transient and receptor desensitization shape synaptic responses evoked by hippocampal neurogliaform cells. The Journal of neuroscience : the official journal of the Society for Neuroscience 30, 9898–9909, doi:10.1523/JNEUROSCI.5883-09.2010 (2010).

51 Letzkus, J. J., Wolff, S. B. & Luthi, A. Disinhibition, a Circuit Mechanism for Associative Learning and Memory. Neuron 88, 264–276, doi:10.1016/j.neuron.2015.09.024 (2015).

52 Kitamura, T. et al. Island cells control temporal association memory. Science 343, 896–901, doi:10.1126/science.1244634 (2014).

53 Basu, J. et al. Gating of hippocampal activity, plasticity, and memory by entorhinal cortex long-range inhibition. Science 351, aaa5694, doi:10.1126/science.aaa5694 (2016).

54 Yamawaki, N. et al. Long-range inhibitory intersection of a retrosplenial thalamocortical circuit by apical tuft-targeting CA1 neurons. Nature neuroscience 22, 618–626, doi:10.1038/s41593-019-0355-x (2019).

55 Fuentealba, P. et al. Expression of COUP-TFII nuclear receptor in restricted GABAergic neuronal populations in the adult rat hippocampus. The Journal of neuroscience : the official journal of the Society for Neuroscience 30, 1595–1609, doi:10.1523/JNEUROSCI.4199-09.2010 (2010).

56 Bezaire, M. J., Raikov, I., Burk, K., Vyas, D. & Soltesz, I. Interneuronal mechanisms of hippocampal theta oscillations in a full-scale model of the rodent CA1 circuit. eLife 5, doi:10.7554/eLife.18566 (2016).

57 Lasztoczi, B. & Klausberger, T. Layer-specific GABAergic control of distinct gamma oscillations in the CA1 hippocampus. Neuron 81, 1126–1139, doi:10.1016/j.neuron.2014.01.021 (2014).

58 Kaifosh, P. & Losonczy, A. Mnemonic Functions for Nonlinear Dendritic Integration in Hippocampal Pyramidal Circuits. Neuron 90, 622–634, doi:10.1016/j.neuron.2016.03.019 (2016).

59 McClelland, J. L. & Goddard, N. H. Considerations arising from a complementary learning systems perspective on hippocampus and neocortex. Hippocampus 6, 654–665, doi:10.1002/(SICI)1098-1063(1996)6:6<654::AID-HIPO8>3.0.CO;2-G (1996).

60 Treves, A. & Rolls, E. T. Computational analysis of the role of the hippocampus in memory. Hippocampus 4, 374–391, doi:10.1002/hipo.450040319 (1994).

61 Whitlock, J. R., Heynen, A. J., Shuler, M. G. & Bear, M. F. Learning induces long-term potentiation in the hippocampus. Science 313, 1093–1097, doi:10.1126/science.1128134 (2006).

62 Nakashiba, T., Young, J. Z., McHugh, T. J., Buhl, D. L. & Tonegawa, S. Transgenic inhibition of synaptic transmission reveals role of CA3 output in hippocampal learning. Science 319, 1260–1264, doi:10.1126/science.1151120 (2008).

63 Lisman, J. E. & Otmakhova, N. A. Storage, recall, and novelty detection of sequences by the hippocampus: elaborating on the SOCRATIC model to account for normal and aberrant effects of dopamine. Hippocampus 11, 551–568, doi:10.1002/hipo.1071 (2001).

64 Dan, Y. & Poo, M. M. Spike timing-dependent plasticity: from synapse to perception. Physiological reviews 86, 1033–1048, doi:10.1152/physrev.00030.2005 (2006).

65 Barbera, G. et al. Spatially Compact Neural Clusters in the Dorsal Striatum Encode Locomotion Relevant Information. Neuron 92, 202–213, doi:10.1016/j.neuron.2016.08.037 (2016).

66 Pnevmatikakis, E. A. & Giovannucci, A. NoRMCorre: An online algorithm for piecewise rigid motion correction of calcium imaging data. Journal of neuroscience methods 291, 83–94, doi:10.1016/j.jneumeth.2017.07.031 (2017).

67 Zhou, P. et al. Efficient and accurate extraction of in vivo calcium signals from microendoscopic video data. eLife 7, doi:10.7554/eLife.28728 (2018).

